# LSD’s effects are differentially modulated in arrestin knockout mice

**DOI:** 10.1101/2021.02.04.429772

**Authors:** Ramona M. Rodriguiz, Vineet Nadkarni, Christopher R. Means, Yi-Ting Chiu, Bryan L. Roth, William C. Wetsel

**Affiliations:** Department of Psychiatry and Behavioral Sciences, Duke University Medical Center, Durham, NC 27710; Mouse Behavioral and Neuroendocrine Analysis Core Facility, Departments of Cell Biology and Neurobiology, Duke University Medical Center, Durham, NC 27710; Department of Pharmacology, Center for Integrative Chemical Biology and Drug Discovery, Division of Chemical Biology and Medicinal Chemistry, Eshelman School of Pharmacy, University of North Carolina at Chapel Hill School of Medicine, Chapel Hill, NC 27599; Department of Pharmacology and National Institute of Mental Health Psychoactive Drug Screening Program, Center for Integrative Chemical Biology and Drug Discovery, Division of Chemical Biology and Medicinal Chemistry, Eshelman School of Pharmacy, University of North Carolina at Chapel Hill School of Medicine, Chapel Hill, NC 27599; Departments of Cell Biology and Neurobiology, Duke University Medical Center, Durham, NC 27710

**Author notes:** Correspondence: Dr. W.C. Wetsel, Department of Psychiatry and Behavioral Sciences, Duke University Medical Center, 354 Sands Building, PO Box 103203, 303 Research Drive, Durham, NC 27710, USA.

## Abstract

Recent evidence suggests that psychedelic drugs can exert beneficial effects on anxiety, depression, and ethanol and nicotine abuse in humans. However, the hallucinogenic side-effects of psychedelics often preclude their clinical use. Lysergic acid diethylamide (LSD) is a prototypical hallucinogen and its psychedelic actions are exerted through the 5-HT_2A_ serotonin receptor (5-HT2AR). 5-HT2AR activation stimulates Gq- and β-arrestin-(βArr) mediated signaling. To separate effects of these signaling modes, we have used βArr1 and βArr2 mice. We find that LSD stimulates motor activities to similar extents in WT and βArr1-KO mice, with non-significant effects in βArr2-KOs. LSD robustly stimulates many surrogates of psychedelic drug actions including head twitches, grooming, retrograde walking, and nose poking in WT and βArr1-KO animals. In contrast, LSD only slightly stimulates head twitches in βArr2-KO mice, without effects on retrograde walking or nose poking. The 5-HT2AR antagonist MDL100907 (MDL) blocks these LSD effects. LSD also disrupts prepulse inhibition (PPI) in WT and βArr1-KOs; PPI is unaffected in βArr2-KOs. MDL restores PPI in WT mice, but this antagonist is without effect and haloperidol is required in βArr1-KOs. LSD produces a biphasic body-temperature response in WT mice, a monophasic response in βArr1-KOs, and is without effect in βArr2 mutants. Both MDL and the 5-HT1AR antagonist, WAY 100635 (WAY), block the effects of LSD on body temperatures in WT mice, whereas WAY is effective in βArr1-KOs. Collectively, these results reveal that LSD produces diverse behavioral effects through βArr1 and βArr2, and that LSD’s psychedelic drug-like actions appear to require βArr2.

## INTRODUCTION

Lysergic acid diethylamide (LSD) is a prototypical psychedelic drug and is recognized as one of the most potent drugs in this class [1]. LSD was synthesized by Albert Hofmann in 1938, who later discovered its hallucinogenic properties [1-2]. LSD alters sensation, perception, thought, mood, sense of time and space, and consciousness of self in humans [1,3]. Since LSD-induced states appear to bear many similarities to early acute phases of psychosis and because serotonin (5-HT) and LSD both contain an indolamine moiety, Woolley and Shaw [4] proposed that aberrant 5-HT levels in brain could produce mental disturbances that included psychosis. This suggestion gave rise to the 5-HT hypothesis for schizophrenia and stimulated researchers to examine LSD responses in the hopes they could gain a better understanding of the basis for schizophrenia. However, this research was largely curtailed when LSD was classified as a DEA Schedule I drug in the 1960’s. Recent research has revealed that LSD has medicinal value in treating cluster headaches [5], anxiety and depressive disorders in life-threatening conditions when combined with psychotherapy [6], and that it may have potential for studying human consciousness and substance abuse [7-8].

As noted above, LSD shares structural similarities to 5-HT [Nichols, 2016]. Thus, it is not surprising that LSD binds to all thirteen of the 5-HT G protein coupled receptors (GPCRs) [9-11]. Besides 5-HT receptors, LSD activates a number of other biogenic amine GPCRs [9] and this polypharmacy may contribute to LSD’s wide ranging activities. One activity in particular regarding LSD has been its hallucinogenic actions. This activity is ascribed to stimulation of the 5-HT 2A receptor (5-HT2AR) since in drug discrimination studies, discrimination potency is highly correlated with hallucinogenic potency in humans [12]. The same psychedelics produce head twitches in mice and, as a result, this response has been proposed as a proxy for hallucinations in humans [13], even though non-psychedelic drugs like 5-hydroxytryptophan induce robust head-twitch responses [14]. Hallucinogen-induced head twitches in rodents are blocked by the highly selective 5-HT2AR antagonist MDL100907 [15-17] and are absent in *htr2A* knockout (KO) mice [18-19]. In addition, recent human studies have shown the hallucinogenic actions of LSD are blocked with the 5-HT2AR preferring antagonist ketanserin [20]. Thus, the hallucinogenic effects of LSD appear to be mediated through the 5-HT2AR [21].

The 5-HT2AR is a member of the rhodopsin family of GPCRs that is coupled to G_q_ protein and β-arrestin (βArr) mediated signaling [22-25]. Recent experiments have found the 5-HT2AR preferentially activates G_q_ family members, with moderate activity at G_z_, and minimal detectible activity at other G_i_, G_12/13_, and G_s_-family members [26]. However, the 5-HT2AR binds to both βArr1 and βArr2 proteins *in vitro* and is complexed with bArr1 and bArr2 in cortical neurons *in vivo* [25]. While most GPCR agonists, like 5-HT, activate both G protein and βArr modes of signaling, it has become established that ligand binding can activate the G protein-dependent signaling pathway while serving to activate or inhibit a G protein-independent pathway through βArr. Hence, a given ligand can act as an agonist at one pathway while inhibiting the other pathway or it can serve various combinations of these actions. This property has been termed functional selectivity or biased signaling [27-29] and ligands have been developed to exploit these signaling features [c.f., 30]. Although LSD activates G protein signaling at many GPCRs [11], this psychedelic stimulates βArr-mediated responses at most tested biogenic amine GPCRs [9]. Interestingly, LSD displays βArr-biased signaling at the 5-HT2AR [10-11,26]. Most 5-HT2AR-containing neurons express both βArr1 and βArr2 [25] and mice have been generated that have *Arrb1* or *Arrb2* deleted [31-32]. The purpose of the present investigations was to determine whether LSD would produce behavioral and physiological effects that were differential among the wild-type (WT), βArr1-KO, and βArr2-KO mice.

## MATERIALS AND METHODS

Details of the materials and methods can be found in the Supplementary Information.

### Subjects

Adult male and female WT and βArr1-KO, and WT and βArr2-KO mice were used in these experiments [1-2]. All studies were conducted with an approved protocol from the Duke University Institutional Animal Care and Use Committee.

### Open field activity

Motor activities were assessed over 120 min in an open field (Omnitech Electronics, Columbus, OH) illuminated at 180 lux [3].

### Head twitch, grooming, and retrograde walking

These behaviors were filmed during the assessment of motor activity in the open field.

### Nose-poking responses

Nose-pokes were monitored in a mouse 5-choice serial reaction-time apparatus (Med Associates Inc., St. Albans, VT) [4].

### Prepulse inhibition (PPI)

PPI of the acoustic startle response was conducted using SR-LAB chambers (San Diego Instruments, San Diego, CA) as reported [3].

### Regulation of body temperature

Baseline body temperatures were taken (Physitemp Instruments LLC, Clifton, NY) in the absence of the vehicle or antagonist. Immediately afterwards mice were injected with vehicle or different doses of MDL or WAY. Fifteen min later (time 0) their body temperatures were taken and they were administered vehicle or LSD. Subsequently, body temperatures were taken at 15, 30, 60, 120, 180, and 240 min.

### Radioligand binding and immunohistochemistry of 5-HT2AR

Radioligand binding experiments on mouse brains were conducted as previously detailed using [^3^H]-ketanserin as the radioligand [5]. The 5-HT2AR immunofluorescence/microscopy was performed as described [35] with a previously validated 5-HT2AR-specific antibody [36].

### Statistics

All statistical analyses were performed with IBM SPSS Statistics 27 programs (IBM, Chicago, IL).

## RESULTS

### Effects of Arrb1 or Arrb2 deletion on LSD-stimulated motor responses

LSD stimulates, inhibits, produces biphasic effects on motor activities in rats [18,37-43]. The βArr1 and βArr2 mice were used to determine whether disruption of either of these gene products could modify responses to LSD and to test whether 5-HT2AR antagonism differentially could block the effects of LSD. Locomotor, rearing, and stereotypical activities were monitored in the βArr1 and βArr2 mice at 5-min intervals over the 120 min test with the βArr1 and βArr2 mice (Supplementary Figs. S1-S2; Supplementary Tables S1-S2). When cumulative baseline locomotion was examined in βArr1 mice, activity was not differentiated by genotype or the pre-assigned treatment condition (Supplementary Table S3; Supplementary Table S4). By contrast, overall cumulative baseline rearing activities were lower in groups that were to receive 0.1 or 0.5 mg/kg MDL with LSD compared to mice that were to be given LSD or the vehicle (*p*-values≤0.016). For stereotypy in βArr1 animals, overall cumulative baseline activities were lower in groups that were to receive 0.1 or 0.5 MDL with LSD than the vehicle control (*p*-values≤0.005); stereotypy in the group to be injected with 0.1 mg/kg MDL with LSD was lower also than the group to be given LSD (*p*=0.002). In comparison to the in βArr1 mice, baseline cumulative locomotor, rearing, and stereotypical activities in the βArr2 animals were not distinguished by genotype or treatment (Supplementary Table S5; Supplementary Table S6)

When cumulative LSD-stimulated motor activities were examined in the βArr1 mice, no genotype effects were found. This psychedelic stimulated locomotion relative to the groups given 0.5 mg/kg MDL alone or the vehicle (*p*-values≤0.001) (Fig. 1a; Supplementary Table S4). Importantly, both 0.1 and 0.5 mg/kg MDL blocked the locomotor-stimulating effects of LSD (*p*-values≤0.001). When cumulative rearing activities were examined, these responses were found to be lower in groups given 0.5 mg/kg MDL alone or administered 0.1 or 0.5 mg/kg MDL with LSD (*p*-values≤0.028) (Fig. 1c). An assessment of cumulative stereotypical activities in the βArr1 animals revealed responses in the MDL group, as well as in the 0.1 and 0.5 mg/kg MDL plus LSD groups to be significantly decreased relative to the LSD-treated mice (*p*-values≤0.027) (Fig. 1e). Hence, MDL normalized the LSD-induced hyperactivity in the βArr1 animals.

**Fig. 1.**
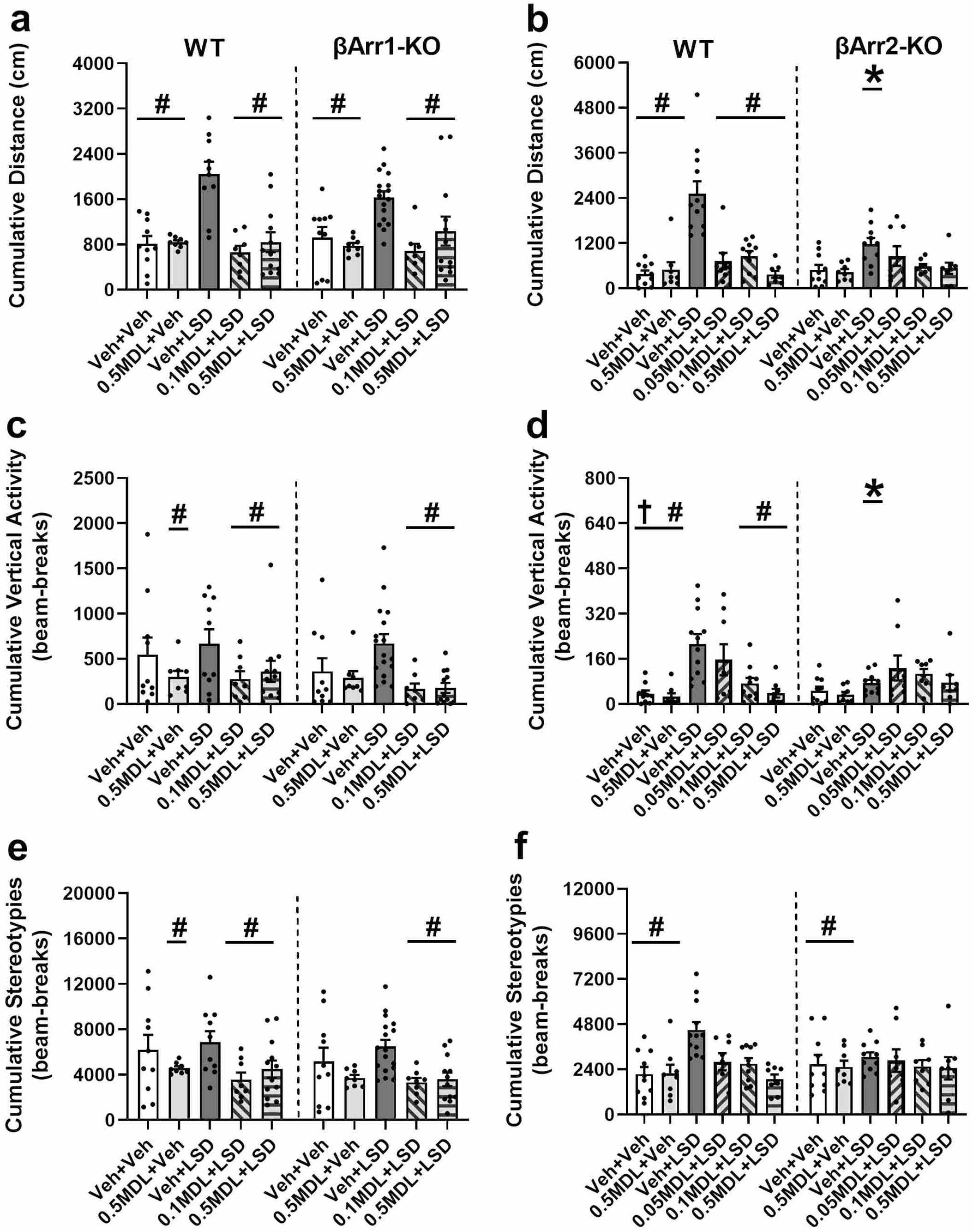
Effects of LSD and MDL100907 on cumulative motor activities in the β-arrestin 1 and β-arrestin 2 mice. Mice were administered the vehicle or different doses of MDL100907 (MDL) and placed into the open field for 30 min. They were removed injected with the vehicle or 0.3 mg/kg LSD and immediately returned to the test arena for 90 min. The cumulative baseline results (0-30 min) are depicted in Supplementary Tables S3 and S5. The statistics for the cumulative LSD-stimulated (31-120 min) motor activities are presented in in Supplementary Tables S4 and S6. **a** Locomotor activities in WT and βArr1-KO mice. **b** Locomotor activities in WT and βArr2-KO subjects. **c** Rearing activities in βArr1 animals. **d** Rearing activities in βArr2 subjects. **e** Stereotypical activities in βArr1 mice. **f** Stereotypical activities in βArr2 animals. N = 8-17 mice/group. ^*^*p*<0.05, WT vs. KO; #*p*<0.05, vs. LSD overall (βArr1 all motor activities; βArr2 for stereotypy) or within genotype (βArr2 for locomotion and rearing); †*p*<0.05, vs. 0.05 mg/kg LSD + LSD.

Effects of LSD in the βArr2-KO mice were quite different from those of its WT controls (Supplementary Table S6). LSD was more potent in stimulating cumulative locomotor and rearing activities in the WT than in βArr2-KO mice (*p*-values<0.001); Fig. 1b,d); no genotype differences were observed for cumulative stereotypical activities (Fig. 1f). When locomotion was analyzed for WT animals, the LSD-stimulated responses were higher than those for all other treatment groups (*p*-values<0.001) (Fig. 1b). Hence, all three doses of MDL were efficacious in suppressing the LSD-induced hyperlocomotion. Although LSD increased locomotor activity in βArr2-KO mice, it was not significantly different from any other treatment group. In WT mice rearing activities were increased with LSD over that for the MDL and vehicle controls (*p*-values<0.001) (Fig. 1d). Although rearing was higher in WT mice given 0.05 mg/kg MDL with LSD than in the MDL and vehicle controls (*p*-values≤0.029), the 0.1 and 0.5 mg/kg doses of MDL reduced the LSD-stimulated rearing activity (*p*-values≤0.001) to the levels of these controls. By comparison, LSD was without effect in the βArr2-KO mice. An examination of stereotypy failed to detect any significant genotype effects (Fig. 1f). Nonetheless, treatment effects were evident overall with LSD stimulating stereotypical activities over that of the vehicle and MDL controls (*p*-values≤0.013); however, when 0.5 mg/kg MDL was given with LSD the psychedelic effects were abrogated (*p*=0.003). Collectively, these results indicate that LSD stimulates motor responses to similar extents in the βArr1 and βArr2 WT mice and in βArr1-KO animals, and that the 5-HT2AR antagonist MDL blocks these LSD-stimulated activities. By striking comparison, LSD exerts minimal effects on these same responses in the βArr2-KO mice where none of these activities were significantly increased above that of the vehicle or MDL controls.

### LSD and MDL100907 effects on additional LSD-stimulated behaviors

LSD affects a number of behaviors in mice [13,18,44-48] that include, as a minimum, head twitches, grooming, and retrograde walking. In addition, LSD produces changes in hole-poking behavior in rats [41-42,45-50]. Despite an absence of genotype effects with βArr1 mice for head twitches, grooming, retrograde walking, and nose-poking, overall treatment effects were evident (Supplementary Table S7). Relative to the MDL and vehicle controls, LSD stimulated head twitches in WT and βArr1-KO mice (*p*-values<0.001) (Fig. 2a). Both 0.1 and 0.5 mg/kg MDL blocked these LSD effects by restoring the numbers of head twitches to those of controls. A similar effect was observed for grooming where LSD augmented this response over that of groups administered the vehicle or 0.5 mg/kg MDL alone (*p*-values<0.001) (Fig. 2c). When 0.1 or 0.5 mg/kg MDL was given with LSD, both doses of the 5-HT2AR antagonist suppressed the LSD stimulatory effect (*p*-values<0.001) to that of the controls.

**Fig. 2.**
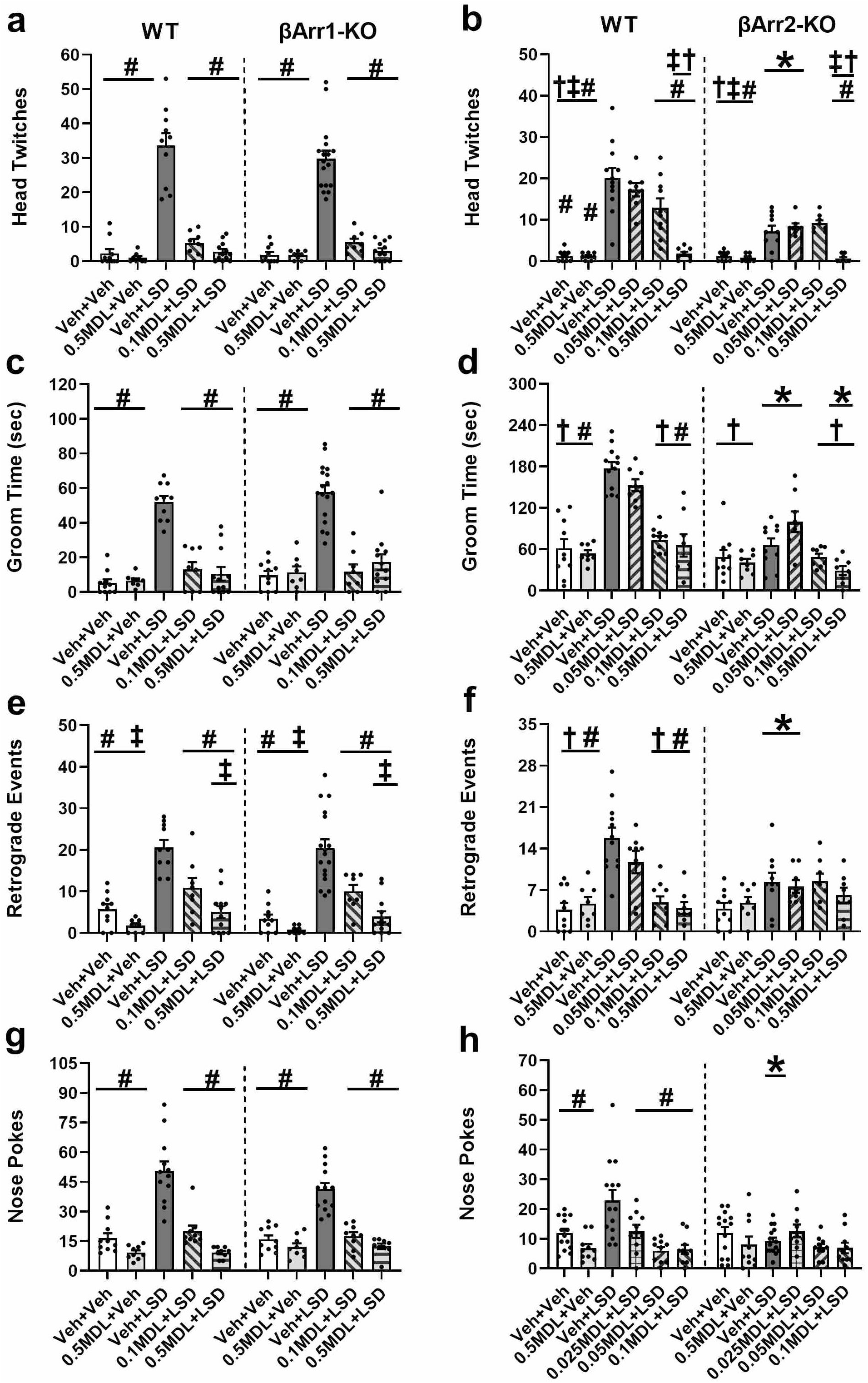
Effects of LSD and MDL100907 on behavioral responses in the β-arrestin 1 and β-arrestin 2 mice. A description of the experimental design is presented in the legend for Figure 1. The head twitch, grooming, and retrograde walking results represent the first 30 min after injection of LSD. Nose poking was examined in a 5-choice serial reaction time apparatus (no rewards) with a similar time-course for vehicle and MDL injections (as in the open field) followed by administration of the vehicle and LSD. The statistics for the βArr1 and βArr2 responses can be found in Supplementary Tables S7 and S8, respectively. **a** Head twitches in WT and βArr1-KO mice. **b** Head twitches in WT and βArr2-KO animals. **c** Duration of grooming in βArr1 animals. **d** Duration of grooming in βArr2 subjects. **e** Incidences of retrograde walking in βArr1 subjects. **f** Incidences of retrograde walking in βArr2 mice. **g** Nose poking in WT and βArr1-KO mice. **h** Nose poking in WT and βArr2-KO animals. N = 8-17 mice/group for head twitch, grooming, and retrograde walking; N = 9-15 mice/group for nose-poking. ^*^*p*<0.05, WT vs. KO; #*p*<0.05, vs. LSD overall (βArr1) or within genotype (βArr2); †*p*<0.05, vs. 0.05 mg/kg MDL + LSD; ‡*p*<0.05, vs. 0.1 mg/kg MDL + LSD overall (βArr1) or within genotype (βArr2).

Besides grooming, LSD was efficacious in potentiating retrograde walking in the βArr1 mice compared to the vehicle and MDL controls (*p*-values<0.001) (Fig. 2e) (Supplementary Table S7). While 0.1 and 0.5 mg/kg MDL depressed this LSD-mediated response (*p*-values<0.001), the lower dose of MDL was less efficacious than the higher dose in normalizing this behavior (*p*=0.010). When nose poking behaviors were examined, LSD was observed to increase this response overall relative to the vehicle and MDL controls (*p*-values<0.001) (Fig. 2g). Both doses of MDL suppressed the LSD stimulation of nose-poking behaviors (*p*-values<0.001).

In contradistinction to the βArr1 mice, genotype differences were found between the βArr2 animals (Supplementary Table S8). The numbers of head twitches were significantly increased in the WT than in βArr2-KO mice treated with LSD alone or 0.05 mg/kg MDL with LSD (*p*-values<0.001) (Fig. 2b). In WT mice head twitches were augmented by LSD and by 0.05 or 0.1 mg/kg MDL plus LSD relative to the MDL and vehicle controls (*p*-values<0.001). While 0.05 mg/kg MDL was ineffective in reducing this LSD-mediated behavior, both the 0.1 and 0.5 mg/kg doses significantly suppressed this response (*p*-values≤0.002) – with the higher MDL dose being the more efficacious (*p*<0.001). The LSD and 0.05 or 0.1 mg/kg MDL plus LSD treatments also increased head twitch behaviors in βArr2-KO animals compared to their MDL and vehicle controls and the 0.5 mg/kg MDL plus LSD treatment (*p*-values≤0.023). Only 0.5 mg/kg MDL was sufficient to depress this LSD-initiated response (*p*=0.019) to the levels of the controls.

For grooming, the durations of grooming were higher in the WT than in the corresponding βArr2-KO groups administered LSD alone, 0.05 mg/kg MDL plus LSD, or 0.5 mg/kg MDL with LSD (*p*-values≤0.016) (Fig. 2d) (Supplementary Table S8). In WT mice LSD augmented grooming relative to the MDL and vehicle controls (*p*<0.001). While 0.05 mg/kg MDL failed to block the LSD effect, both the 0.1 and 0.5 mg/kg doses were efficacious (*p*-values<0.001) in returning responses to control levels. In βArr2-KO animals, LSD failed to increase the duration of grooming responses relative to the MDL and vehicle controls. Nevertheless, grooming was enhanced in the group administered 0.05 mg/kg MDL plus LSD relative to all groups (*p*-values≤0.013), except those given LSD alone.

Aside from disturbing grooming, LSD also induced retrograde walking (Supplementary Table S8). Here, genotype effects were observed in mice that received LSD or 0.05 mg/kg MDL with LSD (*p*-values*≤*0.040) (Fig. 2f). In WT mice LSD potentiated the incidences of retrograde walking compared to the MDL and vehicle controls (*p*<0.001). Although 0.05 mg/kg MDL was ineffective in decreasing this LSD-stimulated behavior, both the 0.1 and 0.5 mg/kg doses suppressed this response (*p*<0.001). By contrast, LSD was without significant effect on retrograde walking in the βArr2-KO animals. Differential nose-poking responses were observed between the βArr2 mice (Fig. 2h). Here, LSD stimulated nose-poking behaviors in WT mice relative to all other groups (*p*<0.001). All doses of MDL suppressed LSD-stimulated nose poking in WT mice (*p*-values≤0.007) to the MDL and vehicle controls. By comparison, nose-poking in βArr2-KO animals was not distinguished by treatment condition.

Since LSD can induce a fragmentation of consciousness [3], we examined grooming in detail since it has a chained organization of responses in rodents [51]. Inspection of the video-recordings confirmed that all genotypes engaged in a normal sequence of grooming beginning with the face, progressing down the body, and ending at the feet or tail (Movie 1). When LSD was administered, the sequence of grooming in the WT and βArr1-KO mice became abbreviated, non-sequential, and/or restricted to one area of the body (Movies 2-3). By sharp comparison, the grooming sequence was complete and rarely perturbed in the βArr2-KO animals (Movie 4). When MDL alone was injected, the organization of grooming was intact in the WT and βArr1-KO mice (Movie 5). With MDL the βArr2-KO animals might pause in the grooming bout and/or display twitching of neck and back muscles but they would finish the sequence (Movie 6). The patterns of grooming among the genotypes with MDL plus LSD were divergent. In WT mice, when MDL was given with LSD, the organization of grooming was restored (Movie 7). When the βArr1 mutants received the same treatment, they began the grooming sequence, engaged in focal grooming of a part of the body, and then completed the sequence (Movie 8). When this same drug combination was administered to βArr2-KO mice, they usually began the sequence appropriately, but at some mid- or later-point they would become focused on one area; however, they usually completed the grooming sequence (Movie 9).

In summary, responses to LSD across these LSD-stimulated behaviors were usually similar between the βArr1 genotypes and MDL reduced these responses to levels of the controls. Importantly, the βArr2 mice responded quite differently than the other genotypes. Responses to LSD were significantly higher overall in WT than in βArr2-KO animals. LSD did not significantly increase grooming, retrograde walking, or nose poking behaviors in these mutants. Notably, LSD disrupted the sequences of grooming in WT mice from both strains and in the βArr1-KO mice; the βArr2-KO animals were unaffected. Nonetheless, divergent responses to MDL alone or MDL plus LSD were observed among the genotypes.

### LSD and MDL100907 effects on prepulse inhibition

LSD disrupts PPI in both rats and humans [44,52] and the response can be restored with a 5-HT2AR antagonist [44,53]. βArr1 mice were pre-treated with the vehicle or with 0.1 or 0.5 mg/kg MDL. Subsequently, they were administered the vehicle or 0.3 mg/kg LSD and tested in PPI. No significant genotype or treatment effects were observed for null activity or in response to the 120 dB startle stimulus (Supplementary Fig. S3a-b; Supplementary Table S9). In contrast, LSD depressed PPI in both βArr1 genotypes (*p*-values≤0.002) relative to the MDL and vehicle controls (Fig. 3a). When WT mice were administered either dose of the 5-HT2AR antagonist with LSD, it restored PPI to control levels. In stark contrast, neither dose of the antagonist was effective in blocking the LSD effects in the βArr1-KO mice. Notably, PPI in these two groups was significantly lower than that in similarly treated WT animals (*p*-values≤0.018).

**Fig. 3.**
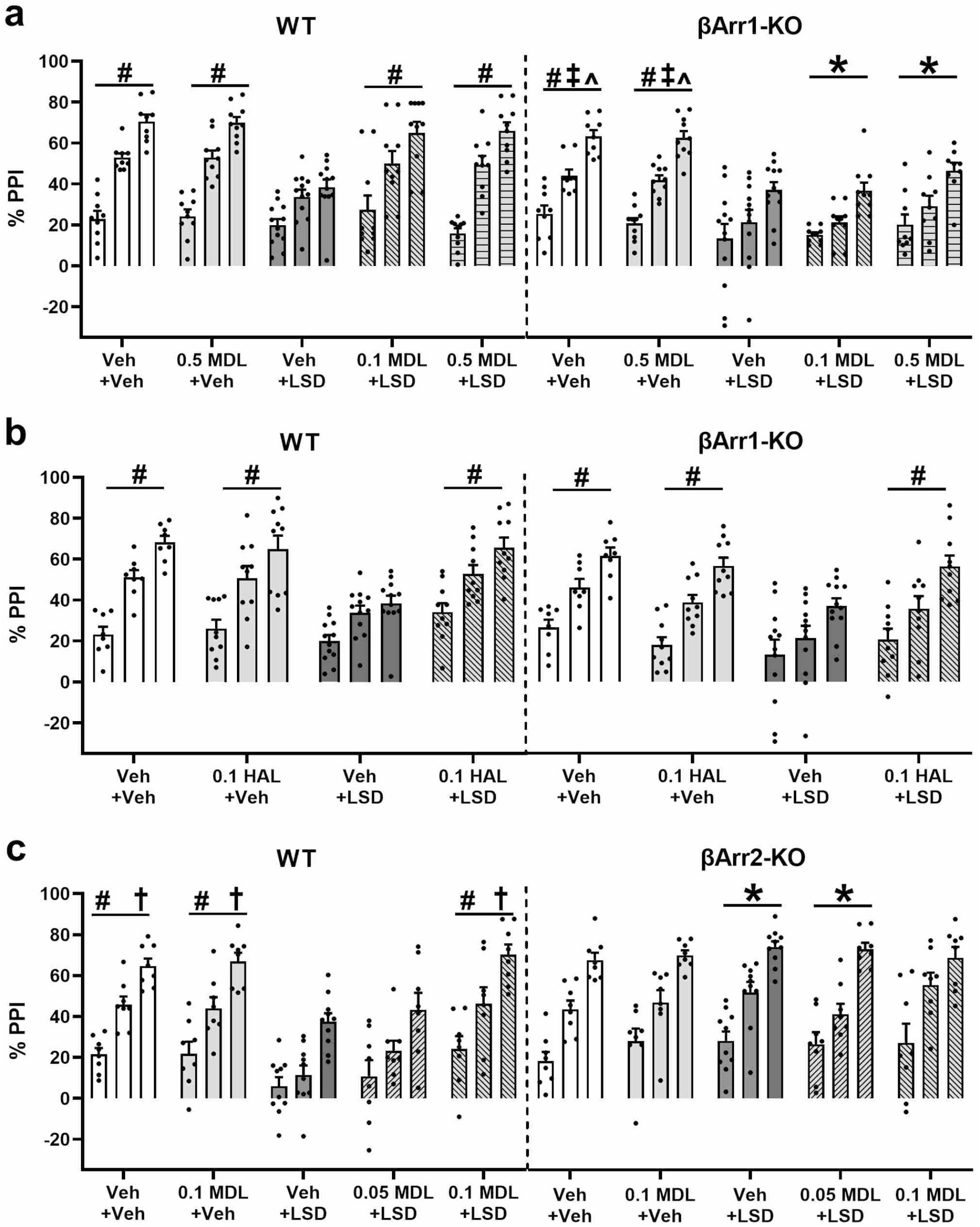
Effects of LSD, MDL100907, and haloperidol on prepulse inhibition in β-arrestin 1 and β-arrestin 2 mice. Mice were injected with MDL100907, haloperidol, or the vehicle and administered subsequently the vehicle or LSD prior to testing PPI. The statistics for the βArr1 and βArr2 responses are located in Supplementary Tables S9-S11. **a** PPI in WT and βArr1-KO mice treated with MDL or LSD. **b** PPI in WT and βArr1-KO mice treated with haloperidol or LSD. **c** PPI in WT and βArr2-KO mice treated with MDL or LSD. N = 8-12 mice/group. ^*^*p*<0.05, WT vs. KO; #*p*<0.05, vs. LSD overall (βArr1 with haloperidol) within genotype (βArr1 and βArr2 with MDL); †*p*<0.05, vs. 0.05 mg/kg MDL + LSD within genotype; ‡*p*<0.05, vs. 0.1 mg/kg MDL + LSD within genotype; ^*p*<0.05, vs. 0.5 MDL + LSD within genotype.

Since haloperidol can normalize PPI in mouse models [33], we tested whether this antipsychotic drug could normalize the LSD-disrupted PPI in the βArr1-KO mice (Supplementary Table S10). Overall null activity was higher in the 0.1 mg/kg haloperidol plus LSD group than in mice treated with haloperidol alone or the vehicle control (*p*-values≤0.009) (Supplementary Fig. S2c). An assessment of startle activity revealed this activity was lower overall in the WT relative to the βArr1-KO mice (*p*=0.028) (Supplementary Fig. S2d). For PPI, overall responses were reduced in the βArr1-KO compared to the WT animals (*p*=0.008; Fig. 3b). Treatment effects were noted where LSD suppressed PPI relative to all other treatment conditions (*p*-values≤0.002). Hence, haloperidol normalized the LSD-disrupted PPI in both βArr1 genotypes.

PPI responses in the βArr2 mice were also examined. Overall null activity was decreased in the combined 0.1 mg/kg MDL plus LSD group compared to the vehicle control and the LSD group (*p*-values≤0.003) (Supplementary Fig. S2e; Supplementary Table S11). No significant differences were detected for startle activity (Supplementary Fig. S2f). Nevertheless, striking genotype differences were evident for PPI (Fig. 3c). Here, responses to LSD and to the 0.05 MDL plus LSD treatment were reduced in WT relative to the βArr2-KO mice (*p*-values≤0.001). In WT animals, LSD suppressed PPI compared to the MDL and vehicle controls (*p*=0.001). The 0.05 mg/kg dose of MDL was insufficient to restore the LSD-disrupted PPI relative to the vehicle and MDL controls, whereas with 0.1 mg/kg MDL normalization occurred. By dramatic comparison, LSD was completely without effect in the βArr2-KO mice. Collectively, these findings show that LSD disrupts PPI in both genotypes of the βArr1 mice and in the βArr2 WT animals. MDL restores PPI in both WT strains, whereby haloperidol is required to normalize it in βArr1-KO mice. By contrast, PPI in βArr2-KO mice is unaffected by this psychedelic.

### LSD and MDL100907 effects on temperature regulation

LSD is reported to increase and to decrease body temperatures in rodents [54] and both the 5-HT2A and the 5-HT-1A receptors are known to mediate changes in temperature [55-56]. Baseline body temperatures were not significantly different between the genotypes (βArr1: WT 37.6 ±0.05 °F, KO 37.6 ±0.10 °F; βArr2: WT 37.7 ±0.10 °F, KO 37.8 ±0.10 °F). Immediately afterwards mice were injected with the vehicle or different doses of MDL or WAY. Fifteen min later (time 0) body temperatures were taken and the mice were administered the vehicle or LSD. Responses to LSD were followed over 4 hours in all genotypes. Initial analyses were run with the vehicle, and MDL or WAY controls and LSD. Subsequent analyses included the vehicle, LSD, and MDL or WAY with LSD.

For βArr1 mice, genotype differences were detected with the vehicle, MDL, and LSD (Supplementary Table S12). Under vehicle conditions, body temperatures were higher in WT mice at 0 and 120 to 240 min compared to the βArr1-KO animals (*p*-values≤0.011) (Fig. 4a-b). With MDL, temperatures in WT animals were lower at 0 min, but were higher at 180 and 240 min than in the mutants (*p*-values≤0.026). LSD produced a biphasic response in WT animals, but only a monophasic effect in mutants with genotype differences at 30, 120, 180, and 240 min (*p*-values≤0.046). Within WT animals, LSD reduced body temperatures at 15 and 30 min, but increased them at 60, 120, and 180 min relative to vehicle (*p*-values≤0.008). By comparison, in βArr1-KO animals LSD only enhanced temperatures at 60 through 240 min (*p*-values≤0.011). Although MDL decreased body temperatures in WT mice at 0, 15, and 30 min (*p*-values≤0.019), it exerted no effects on the mutants at these times.

**Fig. 4.**
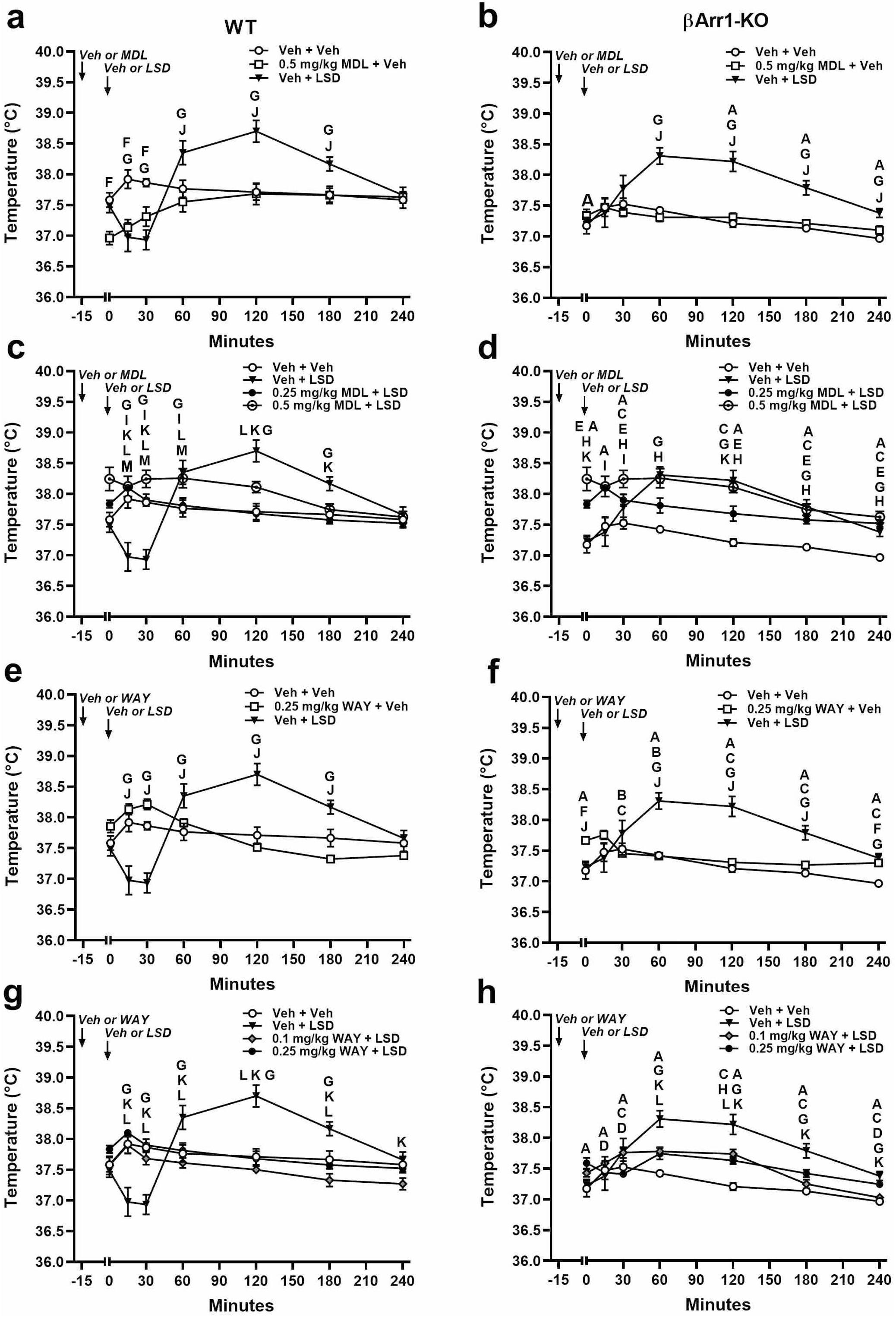
Effects of LSD, MDL100907, and WAY 100635 on body temperature in β-arrestin 1 mice. Mice were injected with the vehicle, MDL, or WAY and 15 min later they received the vehicle or LSD and were examined. The statistics for the βArr1 responses are found in Supplementary Table S12. **a** Body temperatures in WT mice treated with the vehicle, 0.5 mg/kg MDL, or LSD. **b** Temperatures in βArr1-KO animals given similar treatments. **c** Body temperatures in WT mice given the vehicle, LSD, or 0.25 or 0.5 mg/kg MDL plus LSD. **d** Temperatures in similarly treated βArr1-KO animals. **e** Body temperatures in WT mice administered the vehicle, LSD, or 0.25 mg/kg WAY. **f** Temperatures in βArr1-KO animals that received the same treatments. **g** Body temperatures in WT mice that were given the vehicle, LSD, or 0.1 or 0.25 mg/kg WAY with LSD. **h** Temperatures in βArr1-KO animals subjected to the same treatments. N = 9-12 mice/group. ^A^*p*<0.05, WT vs. KO for vehicle; ^B^*p*<0.05, WT vs. KO for low dose MDL or WAY; ^C^*p*<0.05, WT vs. KO for LSD; ^D^*p*<0.05, WT vs. KO for low dose MDL or WAY with LSD; ^E^*p*<0.05, WT vs. KO for high dose MDL or WAY with LSD; ^F^*p*<0.05, vehicle vs. MDL or WAY alone within genotype; ^G^*p*<0.05, vehicle vs. LSD within genotype; ^H^*p*<0.05, vehicle vs. low dose MDL or WAY with LSD within genotype; ^I^*p*<0.05, vehicle vs. high dose MDL or WAY with LSD within genotype; ^J^*p*<0.05, MDL or WAY alone vs. LSD within genotype; ^K^*p*<0.05, LSD vs. low dose MDL or WAY with LSD within genotype; ^L^*p*<0.05, LSD vs. high dose MDL or WAY with LSD within genotype; ^M^*p*<0.05, low dose MDL or WAY with LSD vs. high dose MDL or WAY with LSD within genotype.

Since LSD produced such dramatic effects in body temperatures between the βArr1 genotypes, different doses of MDL were administered with LSD in an attempt to normalize their temperatures (Supplementary Table S12). No genotype differences were obtained when 0.25 mg/kg MDL was given with LSD (Fig. 4c-d). Responses to 0.5 mg/kg MDL with LSD, however, appeared as higher temperatures in WT animals at 0, 30, 120, 180, and 240 min (*p*-values≤0.021) and lower temperatures at 30 min than in the mutants (*p*<0.001). In WT mice, 0.025 mg/kg MDL normalized their LSD-perturbed body temperatures (Fig. 4c). By comparison, this low dose was ineffective in βArr1-KO animals, whereas 0.5 mg/kg MDL was required to normalize their body temperatures to the vehicle control (Fig. 4d).

Because body temperatures in βArr1-KO mice were not fully restored with the 5-HT2AR antagonist, we examined whether a 5-HT1AR antagonist could block LSD’s effects (Supplementary Table S12). Temperature responses to 0.25 mg/kg WAY 100635 (WAY) were significantly higher in WT than mutants at 30 and 60 min (*p*-values≤0.012) (Fig. 4e-f). The LSD-induced responses in WT mice were lower at 30 min but higher from 120 to 240 min than in the βArr1-KO animals (*p*-values≤0.036). In WT mice, body temperatures were lower in WAY-treated mice than in those given LSD at 15 to 180 min (*p*-values≤0.052) (Fig. 4e). By comparison, temperatures in βArr1-KO mice given WAY were higher than the LSD group at 0 min (*p*=0.028), but were lower from 60 min to 180 min (*p*-values≤0.002) (Fig. 4f). Apart from these differences, body temperatures to WAY across time in βArr1-KO mice were different from those of the vehicle group only at 0 and 240 min (*p*-values≤0.046), whereas no significant differences across time were observed for WT mice.

To determine whether WAY exerted any effects on LSD responses, 0.1 or 0.25 mg/kg WAY was administered with this psychedelic (Supplementary Table S12). Although in WT mice both doses of WAY normalized the LSD-induced changes in body temperatures to the vehicle control, 0.25 mg/kg WAY appeared to be the more efficacious (Fig. 4g). While there were some variabilities across time in mutants with the 0.1 and 0.25 mg/kg doses, their LSD-disturbed temperatures were restored to levels that were statistically indistinguishable from the vehicle— with the exception of 0.1 mg/kg WAY at 120 min (*p*=0.011) (Fig. 4h). Together, 0.25 mg/kg MDL and 0.25 mg/kg WAY were most efficacious in WT mice, while 0.5 mg/kg MDL was partially effective at later times and 0.25 mg/kg WAY was most proficient in the mutants.

Effects of LSD on temperature regulation were examined also in the βArr2 mice (Supplementary Table S13). Responses to the vehicle or 0.5 mg/kg MDL were undifferentiated by genotype (Fig. 5a-b). By comparison, the LSD-induced changes in body temperatures were lower at 15 min (*p*=0.001) and higher at 60, 120, and 180 min in WT than in the βArr2-KO mice (*p*-values≤0.019). In WT animals, LSD produced biphasic changes in temperatures at 30, 120, and 180 min relative to vehicle (*p*-values≤0.032) (Fig. 5a). By contrast, no significant LSD effects were observed in βArr2-KO mice at any time-point (Fig. 5b). Temperature responses to MDL in mutants, however, were decreased relative to vehicle at 15 and 60 min (*p*-values≤0.013). When 0.25 mg/kg MDL was given with LSD, no differential genotype effects were noted (Fig. 5c-d). Nonetheless, body temperatures in WT mice administered 0.5 mg/kg MDL plus LSD were higher than in the mutants (*p*-values≤0.032). This same dose also produced deviations from the vehicle control in WT animals at 15 and 30 min (*p*-values≤0.035) and in βArr2-KO mice at 15, 60, and 120 min (*p*-values≤0.035). Regardless, in WT animals 0.025 mg/kg MDL normalized the LSD-induced changes in body temperature across time (Fig. 5c).

**Fig. 5.**
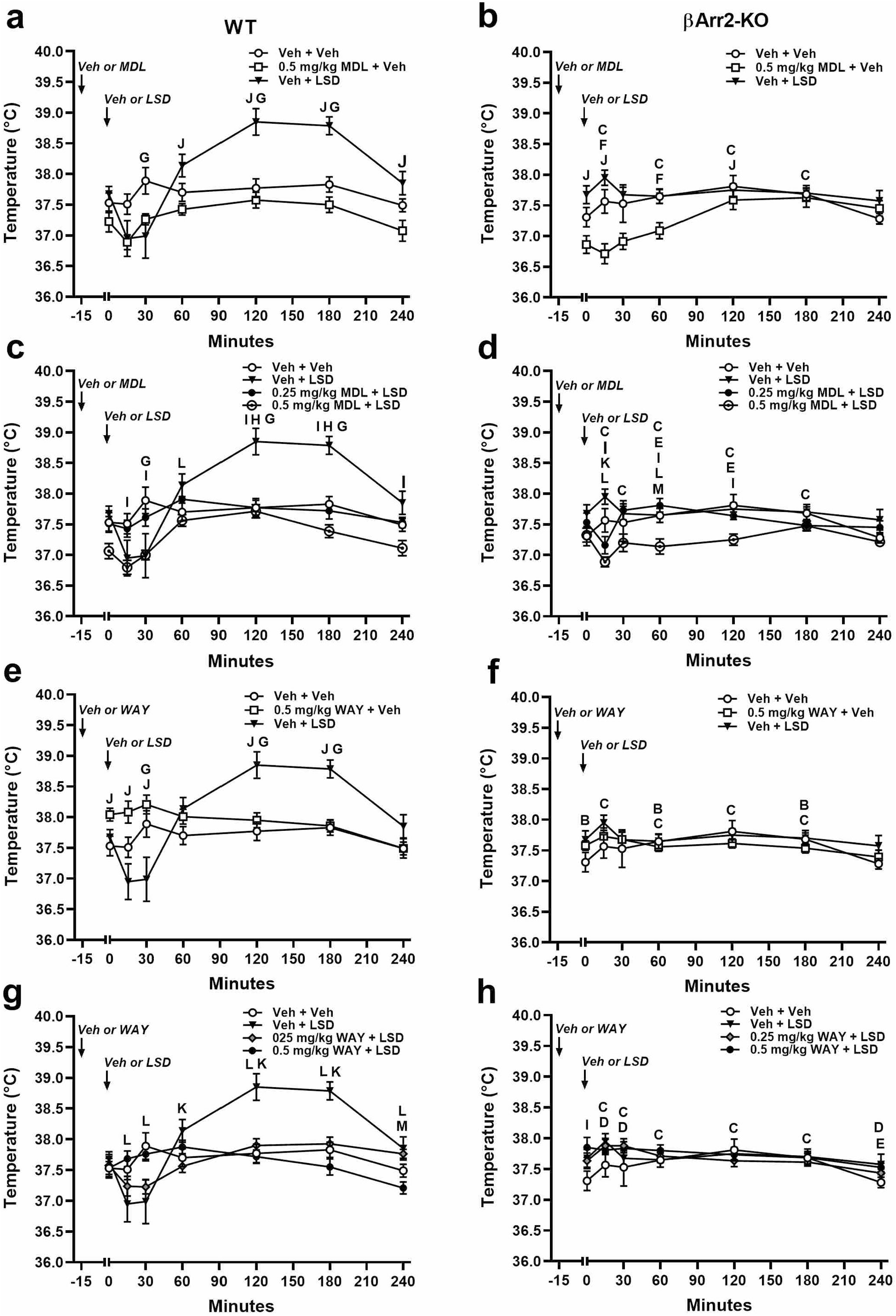
Effects of LSD, MDL100907, and WAY 100635 on body temperature in β-arrestin 2 mice. The statistics for the βArr2 responses are located in Supplementary Table S13. **a** Body temperatures in WT mice treated with the vehicle, 0.5 mg/kg MDL, or LSD. **b** Temperatures in βArr2-KO animals given similar treatments. **c** Body temperatures in WT mice given the vehicle, LSD, or 0.25 or 0.5 mg/kg MDL plus LSD. **d** Temperatures in similarly treated and βArr2-KO animals. **e** Body temperatures in WT mice administered the vehicle, LSD, or 0.25 mg/kg WAY. **f** Temperatures in βArr2-KO animals that received the same treatments. **g** Body temperatures in WT mice that were given the vehicle, LSD, or 0.1 or 0.25 mg/kg WAY with LSD. **h** Temperatures in βArr2-KO animals subjected to the same treatments. N = 8-14 mice/group. ^B^*p*<0.05, WT vs. KO for low dose MDL or WAY; ^C^*p*<0.05, WT vs. KO for LSD; ^D^*p*<0.05, WT vs. KO for low dose MDL or WAY with LSD; ^E^*p*<0.05, WT vs. KO for high dose MDL or WAY with LSD; ^F^*p*<0.05, vehicle vs. MDL or WAY alone within genotype; ^G^*p*<0.05, vehicle vs. LSD within genotype; ^H^*p*<0.05, vehicle vs. low dose MDL or WAY with LSD within genotype; ^I^*p*<0.05, vehicle vs. high dose MDL or WAY with LSD within genotype; ^J^*p*<0.05, MDL or WAY alone vs. LSD within genotype; ^K^*p*<0.05, LSD vs. low dose MDL or WAY with LSD within genotype; ^L^*p*<0.05, LSD vs. high dose MDL or WAY with LSD within genotype; ^M^*p*<0.05, low dose MDL or WAY with LSD vs. high dose MDL or WAY with LSD within genotype.

Effects of WAY were evaluated next in the βArr2 mice (Supplementary Table S13). Genotype differences in response to the 0.5 mg/kg WAY control were observed at 0, 60, and 180 min (*p*-values≤0.023) (Fig. 5e-f). Body temperatures were modulated by LSD in WT relative to βArr2-KO animals at 15, 60, 120, and 180 min (*p*-values≤0.017). Within WT mice, temperatures were enhanced with WAY at 0 min relative to the vehicle control (*p*=0.021) (Fig. 5e). In contrast, responses to WAY were not evident in βArr2-KO animals (Fig. 5f). When 0.25 or 0.5 mg/kg WAY was administered with LSD, genotype differences were found at 15, 30, and 240 min with the lower dose (*p*-values≤0.048) and at 240 min with the higher dose (*p*=0.024) (Fig. 5g-h). For WT mice, 0.25 or 0.5 mg/kg WAY restored the LSD-disturbed body temperatures to levels that were not significantly different from the vehicle control (Fig. 5g). In summary, the low doses of MDL and WAY were both efficacious in reverting the LSD-induced changes to normal in WT animals. By comparison, LSD was without effect in βArr2-KO mice.

### Effects of Arrb1 or Arrb2 deletion on 5-HT2AR expression

Finally, we examined whether deletion of *Arrb* could alter 5-HT2AR expression by radioligand binding with brains from WT and βArr1-KO, and WT and βArr2-KO littermates. When [^3^H]-ketanserin affinity binding was examined, it was found to be significantly higher in βArr2-KO samples than in βArr1 WT and KO brains (*p*-values≤0.039) (Supplementary Fig. S4a). By contrast, the numbers of 5-HT2AR binding sites were higher in brain extracts from βArr2-KO mice than from βArr1 WT and KO animals and the βArr2 WT mice (*p*-values≤0.0.34); no significant differences were observed among the latter three genotypes (Supplementary Fig. S4b). We examined also 5-HT2AR immunofluorescence in βArr1 and βArr2 brain sections (Supplementary Fig. S4c-f). Here, we detected no apparent alterations in the relative receptor distributions among the genotypes. Together, these results are consistent with the hypothesis that neither *Arrb1* nor *Arrb2* genetic deletion *<u>decreases</u>* 5-HT2A receptor expression. Nonetheless, *Arrb2* disruption leads to increased expression of 5-HT2ARs in brain.

## DISCUSSION

In the present study, we analyzed whether global deletion of *Arrb1* or *Arrb2* modifies responses to LSD in the mutants relative to their respective WT controls. In the open field 0.3 mg/kg LSD stimulated motor activities in the βArr1 and βArr2 mice. While these activities were observed in WT animals from both βArr strains and in the βArr1-KO mice. LSD failed to modify significantly rearing or stereotypical activities in WT and βArr1-KO mice. By comparison, LSD stimulated locomotor and rearing activities in βArr2 WT animals, while LSD was without effect in the βArr2-KO mice. Together, these results indicate that LSD-induced motor activities are regulated primarily through βArr2-mediated signaling. In this regard, βArr2 has been reported to play a similar role in morphine-stimulated hyperlocomotion [57] and amphetamine-stimulated locomotor and rearing activities in βArr2 mice [58].

In our experiments, LSD augmented motor activities in βArr1 and βArr2 WT mice and in βArr1-KO animals. By contrast, in rats this hallucinogen has been reported to inhibit ambulation and rearing [40], to stimulate locomotor activities [37,43], to induce divergent effects on motor responses [38], or to produces biphasic (i.e., inhibitory, then stimulatory) effects [39,41-42]. In addition, a sex effect in rats has been observed with LSD [46]. In our experiments with mice, we failed to discern LSD effects attributable to sex. The inhibitory effects of LSD in rats were seen very soon after placement or entry into the open field. In our open field experiments, we did not observe any inhibition of responses with 0.3 mg/kg LSD; only stimulatory effects were evident. This absence of LSD-induced inhibitory effects in our studies could be attributed to differences in the species tested, the dose of LSD used, the test environment and apparatus, and/or the test procedure. For instance, in humans LSD’s behavioral effects are well-known to be context specific [1-3] and the 30 min habituation to the novel environment prior to LSD injection may have reduced emotionality in our mice, such that only the stimulatory effects of LSD were evident.

To determine whether the motor-stimulating effects of LSD were due to 5-HT2AR activation, MDL was used as an antagonist. When administered alone, 0.5 mg/kg MDL exerted no effects on motor performance in either βArr mouse strain. Parenthetically, MDL has been found also to be without effect on spontaneous motor activity in rats [59]. Nonetheless, 0.1 and 0.5 mg/kg MDL blocked the motor-stimulating effects of LSD in the βArr1 and βArr2 WT mice and in the βArr1-KO animals. A similar effect has been attributed to MDL’s effects on LSD-stimulated motor activities in rats [43]. Hence, at the doses tested in our experiments, the present results indicate that the LSD-induced hyperactivity in the βArr mice is promoted through the 5-HT2AR.

Besides motor activity, we examined the effects of LSD on head twitch, grooming, retrograde walking, and nose-poking behaviors. LSD and other psychedelics are well-known to stimulate head-twitch responses in mice [18, 45,48] and this response has been proposed as a proxy for hallucinations in humans [13]. Compared to the vehicle, LSD stimulated head-twitch responses to similar extents in WT and βArr1-KO mice. This psychedelic also activated head twitches in βArr2 animals; however, responses in WT controls were significantly more robust than in βArr2-KO mice. It should be emphasized that these results were surprising since the numbers of 5-HT2AR binding sites were increased in brains of the βArr2-KO animals relative to the other genotypes and because head twitches are mediated through the this receptor [18]. Regardless, in both βArr1 and βArr2 mice, MDL reduced the numbers of head twitches to levels of the vehicle controls. These findings are consistent not only with the known action of MDL on blocking head-twitch responses to various hallucinogens [15-17], but also to the inability of LSD and other psychedelics to induce this response in the *htr2A* homozygous mutant mice [18-19,45].

Aside from head twitches in rodents, LSD stimulates grooming behaviors in cats [60] and it can stimulate or inhibit grooming in mice [46-47]. In our investigations, LSD augmented grooming in βArr1 and βArr2 WT mice, and in βArr1-KO animals. By comparison, LSD failed to prolong grooming in βArr2-KO mice beyond that of vehicle controls. In all cases, 0.1 and 0.5 mg/kg MDL returned the LSD-stimulated grooming to levels indistinguishable from the controls. Thus, antagonism of the 5-HTR2A was sufficient to restore LSD-induced grooming to baseline levels.

Effects of LSD were examined also on the organization of grooming. Under vehicle treatment, all mice displayed similar patterns of grooming that began with the face, progressed to the flanks, and ended with the feet or tail. LSD disorganized this sequence of events such that in the WT and βArr1-KO mice, grooming was often restricted to the flanks, feet, or tail, was non-sequential, or was fragmented. By comparison, grooming in the βArr2-KO mice was largely unaffected by LSD. MDL did not alter grooming in the WT and βArr1-KO mice, whereas it prolonged grooming and promoted twitching of the neck and back muscles in the βArr2 mutants. This antagonist blocked the LSD-disrupting effects on grooming in WT mice and it mostly restored the organization of grooming in βArr1-KO animals. The MDL-LSD combination in βArr2-KO animals produced some disturbances, but the mice typically completed the grooming sequence. Together, these results suggest that additional receptor systems may be involved in these LSD-induced grooming responses. It should be emphasized that LSD perturbs grooming also in rats [43] and cats [60].

LSD effects on retrograde walking and nose-poking responses were examined in the βArr mice. Retrograde walking was one of the first LSD-elicited responses identified in mice [442]. In the behavioral pattern monitor, LSD decreased and then increased or maintained the frequencies of head-poking behaviors on a hole-board [41-42,49-50]. Instead of head-poking on a hole-board, we have used the 5-choice serial reaction-time apparatus to evaluate effects of LSD on nose-poking behavior. LSD stimulated retrograde walking and nose-poking behaviors in WT animals from both βArr strains as well as in the βArr1-KO mice. However, LSD promoted neither response in βArr2-KO animals. Nevertheless, in the other genotypes MDL restored retrograde walking and nose-poking to the levels of vehicle controls. Hence, this 5-HT2AR antagonist normalized these LSD-stimulated responses.

LSD-induced states share many similarities with the early acute phases of psychosis [3]. PPI is abnormal in individuals diagnosed with schizophrenia [61] and LSD disrupts PPI in rats [43,46,53]. In βArr1 mice LSD impaired PPI in both genotypes without affecting startle or null activities. Both 0.1 and 0.5 mg/kg MDL restored LSD-disrupted PPI, but only in the WT mice; an effect consistent with the action of the 5-HT2AR antagonist MDL11,939 in rats [53]. By comparison, βArr1-KO animals were unresponsive to MDL. Since LSD activates human dopamine D2 receptors [9,62], we used haloperidol as a D2 antagonist. At a dose of 0.1 mg/kg, haloperidol restored the LSD-disrupted PPI in βArr1-KO mice. Parenthetically, both 0.1 and 0.2 mg/kg haloperidol failed to rescue PPI in rats given 0.1 mg/kg LSD (s.c.) [43]; the possible reasons for this discrepancy in mice versus rats are unclear. When βArr2 mice were tested, LSD was observed to impair PPI only in WT mice. Notably, βArr2-KO mice were completely unresponsive to this psychedelic. As with βArr1 WT animals, 0.5 mg/kg MDL normalized the LSD-disrupted PPI in βArr2 WT animals. Thus, the LSD effects on PPI in the βArr mice are complex, with restoration of PPI with MDL in both strains of WT mice, normalization of PPI with haloperidol in βArr1-KO animals, and without any discernable effect in the βArr2-KO subjects.

Aside from behavioral studies, LSD effects on body temperatures were evaluated. Parenthetically, LSD serves as an agonist at both 5-HT1ARs and 5-HT2ARs [10-11], and stimulation of either receptor can modify body temperatures in rats [63]. LSD is reported to increase or have no effects on temperatures in mice and to increase or decrease temperatures in rats [54]. We found LSD to alter body temperatures in a biphasic fashion in βArr1 and βArr2 WT mice, in a monophasic manner in βArr1-KO animals, and to be without effect in βArr2-KO subjects. Both MDL and WAY normalized these LSD effects in both WT strains and in βArr1-KO mice. Neither antagonist appeared to be more efficacious than the other. This finding should not be surprising since functional interactions have been described between the 5-HT1ARs and 5-HT2ARs in regulating body temperatures [61]. Although we did not test it, 5-HT2CR antagonists are known to participate in this process. Nevertheless, it appears that antagonism of 5-HT1ARs and 5-HT2ARs is sufficient to normalize the LSD-induced changes in body temperatures in βArr1 and βArr2 mice.

LSD and other psychedelics are well-known for their hallucinogenic actions [1] and these responses have been attributed to 5-HT2AR agonism [12]. We observed LSD to stimulate motor activity, head twitches, grooming, retrograde walking, and nose-poking in the βArr1 and βArr2 WT mice and in βArr1-KO animals. LSD also disrupted PPI and produced diverse changes in body temperatures in these same mice. The LSD-elicited responses in βArr2-KO mice were either significantly attenuated or completely absent. In conditions where LSD produced changes in behavior, these alterations were blocked with the 5-HT2AR antagonist MDL. While these results suggest that the 5-HT2AR is an essential component for all these responses, it should be recalled that LSD exerts a plethora of actions at many GPCRs [9-11] and, aside from head twitch, other responses are inconsistently affected by hallucinogens [18]. Hence, it is likely that LSD’s effects on the 5-HT2AR are involved in a cascade of many GPCR-signaling events mediating these varied responses.

Like other GPCRs, agonist actions at the 5-HT2AR can lead to G protein-dependent and –independent signaling, the latter involves βArr [23-25]. While both βArr1 and βArr2 are ubiquitously expressed in adult rodent brain, expression of βArr2 mRNA is much higher than that for βArr1 – except in selected brain areas [64]. Thus, it may not be surprising that the LSD-elicited responses were less disturbed in the βArr2-KO than in the βArr1-KO mice, because the latter is still retained intact βArr2-mediated signaling. In this regard, it was especially intriguing that LSD-induced head twitch responses were much more robust in the βArr1 and βArr2 WT mice and in the βArr1-KO animals, than in the βArr2 mutants. Our results with LSD indicate that βArr2 may be essential for the expression of hallucinogenic-like actions. Hence, it may be possible to develop novel functionally-selective ligands that preferentially signal though G protein at the 5-HT2AR and avoid the hallucinogenic effects modulated through βArr.

## ACKNOWLEDGEMENTS

We thank Dr. Robert J. Lefkowitz (Duke University Medical Center, Durham, NC, USA) for providing us with the new strain of βArr1 mice and Dr. Laura M. Bohn (Scripps Research Institute, Jupiter, FL, USA) for sending us the βArr2 mice. We thank Mr. Mitchell Huffstickler for the representative videos of grooming behaviors in mice and Ms. Jiechun Zhou for breeding, genotyping, and maintaining the βArr1 and βArr2 mice. Some of the behavioral experiments were conducted with equipment and software purchased with a North Carolina Biotechnology Center grant.

## AUTHOR CONTRIBUTIONS

Mr. Vineet Nadkarni conducted initial open field studies and some ethological experiments with the βArr2 mice and helped to statistically analyze these preliminary data. Mr. Christopher Means conducted the behavioral experiments with the βArr1 and βArr2 mice and organized the data. Dr. Ramona M. Rodriguiz oversaw the experiments, conducted the temperature regulation studies, statistically analyzed the data, and graphed the results. Dr. Yi-Ting Chiu conducted the radioligand binding and immunohistochemical investigations with the βArr1 and βArr2 mice. Drs. Wetsel and Roth conceived the experiments, proposed the experimental designs, and wrote the manuscript.

## REFERENCES

1. Nichols DE. Psychedelics. Pharmacol Rev 2016;68:264–355.

2. Hofmann A. How LSD originated. J Psychedelic Drugs 1979;11:53–60.

3. Geyer MA, Vollenweider FX. Serotonin research: contributions to understanding psychoses. Trends Pharmacol Sci 2008;29:445–53.

4. Woolley DW, Shaw E. A biochemical and pharmacological suggestion about certain mental disorders. Science 1954;119:587–8.

5. Sewell RA, Halpern JH, Pope HG Jr. Response to cluster headache to psilocybin and LSD. Neurology 2006;66:1920–2.

6. Gasser P, Kirchner K, Passie T. LSD-assisted psychotherapy for anxiety associated with a life threatening disease: A qualitative study of acute and sustained subjective effects. J Psychopharmacol 2015;29:57–68.

7. Bogenschutz MP, Johnson MW. Classic hallucinogens in the treatment of addictions. Prog Neuropsychopharmacol Biol Psychiatry 2016;64:250–8.

8. Carhart-Harris RL, Muthukumaraswamy S, Roseman L, Kaelen M, Droog W, Murphy K, et al. Neural correlates of the LSD experience revealed by multimodal neuroimaging. Proc Natl Acad Sci USA 2016;113:4853–8.

9. Kroeze WK, Sassano MF, Huang X-P, Lansu K, McCorvy JD, Giguère PM, et al. PRESTO-Tango as an open source resource for interrogation of the druggable human GPCRome. Nat Struct Mol Biol 2015;22:362–9.

10. Wacker D, Wang C, Katritch V, Han GW, Huang XP, Vardy E, et al. Structural features for functional selectivity at serotonin receptors. Science 2013;340:615–9.

11. Wang C, Jiang Y, Ma J, Wu H, Wacker D, Katritch V, et al. Structural basis for molecular recognition at serotonin receptors. Science 2013;340:610–4.

12. Glennon R. Do classical hallucinogens act as 5-HT_2_agonists or antagonists? Neuropsychopharmacology 1990;3:509–517.

13. Corne SJ, Pickering RW. A possible correlation between drug-induced hallucinations in man and a behavioral response in mice. Psychopharmacologia (Berl.) 1967;11:65–78.

14. Malick JB, Doren E, Barnett A. Quipizine-induce head-twitch in mice. Pharmacol Biochem Behav 1977;6:325–329

15. Fantegrossi WE, Harrington AW, Eckler JR, Arshad S, Rabin RA, Winter JC, et al. Hallucinogen-like actions of 2,5-dimethoxy-4-(n)-propylthiophenethylamine *2C-T-7) in mice and rats. Psychopharmacology (Berl.) 2005;181:496–503.

16. Fantegrossi WE, Harrington AW, Kiessel CL, Eckler JR, Rabin RA, Winter JC, et al. Hallucinogen-like actions of 5-methoxy-N,N-diisopropyltryptamine in mice and rats. Pharmacol Biochem Behav 2006;83:122–9.

17. Fantegrossi WE, Reissig CJ, Katz EB, Yarosh HL, Rice KC, Winter JC. Hallucinogen-like effects of N,N-dipropyltryptamine (DPT): possible mediation by serotonin 5-HT_1A_and 5-HT_2A_receptors in rodents. Pharmacol Biochem Behav 2008;88:358–65.

18. González-Maeso J, Weisstaub NV, Zhou M, Chan P, Ivic L, Ang R, et al. Hallucinogens recruit specific cortical 5-HT_2A_receptor-mediated signaling pathways to affect behavior. Neuron 2007;53;439–52.

19. Keiser MJ, Setola V, Irwin JJ, Laggner C, Abbas AI, Hufeisen SJ, et al. Predicting new molecular targets for known drugs. Nature 2009;462:175–81.

20. Preller KH, Razi A, Zeidman P, Stämpfli P, Friston KJ, Vollenweider FX. Effective connectivity changes in LSD-induced altered states of consciousness in humans. Proc Natl Acad Sci USA 2019;116:2743–48.

21. Roth BL, Willins DL, Kristiansen K., Kroeze WK. Activation is hallucinogenic and antagonism is therapeutic: role 5-HT_2A_receptors in antipsychotic drug actions. The Neuroscientist 1999;5:254–262.

22. Roth BL, Nakaki T, Chuang DM, Costa E. Aortic recognition sites for serotonin (5HT) are coupled to phospholipase C and modulate phosphatidylinositol turnover. Neuropharmacology 1984;23:1223–5.

23. de Chaffoy de Courcelles D, Leysen J E, De Clerck F, Van Belle H, and Janssen PA. Evidence that phospholipid turnover is the signal transducing system coupled to serotonin-S_2_receptor. J Biol Chem 1985;260:7603–7608.

24. Roth BL, Nakaki T, Chuang DM, Costa E. 5-Hydroxytryptamine_2_receptors coupled to phospholipase C in rat aorta: modulation of phosphoinositide turnover by phorbol ester. J Pharmacol Exp Ther 1986;238:480–485.

25. Gelber EI, Kroeze WK, Willins DL, Gray JA, Sinar CA, Hyde EG, et al. Structure and function of the third intracellular loop of the 5-hydroxytryptamine_2A_receptor: the third intracellular loop is α-helical and binds purified arrestins. J Neurochem 1999;72: 2006–14.

26. Kim K, Che T, Panova O, DiBerto JF, Lyu J, Krumm BE, et al. Structure of a hallucinogen-activated Gq-coupled 5-HT_2A_serotonin receptor. Cell 2020;182:1574–1588.

27. Gay EA, Urban JD, Nichols DE, Oxford GS, Mailman RB. Functional selectivity of D_2_receptor ligands in a Chinese hamster ovary hD_2L_cell line: evidence for induction of ligand-specific receptor states. Mol Pharmacol 2004;66:97–105.

28. Urban JD, Clarke WP, von Zastrow M, Nichols DE, Kobilka B, Weinstein H, et al. Functional selectivity and classical concepts of quantitative pharmacology. J Pharm Exp Ther 2007;320:1–13.

29. Violin JD, Lefkowitz RJ. β-arrestin-biased ligands at seven-transmembrane receptors. Trends Pharmacol Sci 2007;28:416–22.

30. Allen JA, Yost JM, Setola V, Chen X, Sassano MF, Chen M, et al. Discovery of β-arrestin-biased D_2_ligands for probing signal transduction pathways essential for antipsychotic efficacy. Proc Natl Acad Sci USA 2011;108:18488–93.

31. Bohn LM, Lefkowitz RJ, Gainetdinov RR, Peppel K, Caron MG, Lin FT. Enhanced morphine analgesia in mice lacking β-arrestin 2. Science 1999;286:2495–8.

32. Kim J, Grotegut CA, Wisler JW, Li T, Mao L, Chen M, et al. β-Arrestin 1 regulates β2-adrenergic receptor-mediated skeletal muscle hypertrophy and contractility. Skelet Muscle 2018;8:39.

33. Park SM, Chen M, Schmerberg C, Dulman R, Rodriguiz RM, Caron MG, et al. Effects of β-arrestin-biased dopamine D2 receptor ligands on schizophrenia-like behavior in hypoglutamatergic mice. Neuropsychopharmacology 2016;41:704-15.

34. Velagapudi R, Subramaniyan S, Xiong C, Porkka F, Rodriguiz RM, Wetsel WC, et al. Orthopedic surgery triggers attention deficits in a delirium-like mouse model. Front Immunol 2019;10: 2675.

35. Yadav PN, Kroeze WK, Farrell MS, Roth BL. Agonist functional selectivity: 5-HT_2A_serotonin receptor antagonist differentially regulate 5-HT_2A_protein level in vivo. J Pharmacol Exp Ther 2011;339:99–105.

36. Magalhaes AC, Holmes KD, Dale LB, Comps-Agrar L, Lee D, Yadav PN, et al. Crf receptor I regulates anxiety behavior via sensitization of 5-HT2 receptor signaling. Nat Neurosci 2011;13:622–9.

37. Dandiya PC, Gupta BD, Gupta ML, Patni SK. Effects of LSD on open field performance in rats. Psychopharmacologia (Berl.) 1969;15:333–40.

38. Gupta BD, Dandiya PC, Gupta ML, Gabba AK. An examination of the effect of central nervous stimulant and anti-depressant drugs on open filed performance in rats. Eur J Pharmacol 1971;13:341–6.

39. Kabeš J, Fink Z, Roth Z. A new device for measuring spontaneous motor activity – effects of lysergic acid diethylamide in rats. Psychopharmacologia (Berl.) 1972;23:75–85.

40. Hughes RN. Effects of LSD on exploratory behavior and locomotion in rats. Behav Biol 1973;9:357–65.

41. Adams LM, Geyer MA. LSD-induced alterations of locomotor patterns and exploration in rats. Psychopharmacology 1982;77:179–185.

42. Mittman SM, Geyer MA. Dissociation of multiple effects of acute LSD on exploratory behavior in rats by ritanserin and propranolol. Psychopharmacology 1991;105:69–76.

43. Ouagazzal A, Grottick AJ, Moreau J, Higgins GA. Effect of LSD on prepulse inhibition and spontaneous behavior in the rat. Neuropsychopharmacology 2001;25:565–75.

44. Woolley DW. Production of abnormal (psychotic?) behavior in mice with lysergic acid diethylamide, and its partial prevention with cholinergic drugs and serotonin. Proc Natl Acad Sci USA 1955;41:338–44.

45. González-Maeso J, Yuen T, Ebersole BJ, Wurmbach E, Lira A, Zhou M, et al. Transcriptome fingerprints distinguish hallucinogenic and nonhallucinogenic 5-hydroxytryptamine 2A receptor agonist effects in mouse somatosensory cortex. J Neurosci 2003;23:8836–43.

46. Páleníček T, Hliňák Z, Bubeníková-Valešová V, Novák T, Horáček J. Sex differences in the effects of N,N-diethyllysergamide (LSD) on behavioural activity and prepulse inhibition. Prog Neuropsychopharmacol Biol Psychiatry 2010;34:588–96.

47. Kyzar EJ, Stewart AM, Kalueff AV. Effects of LSD on grooming behavior in serotonin transporter heterozygous (Sert^+/-^) mice. Behav Brain Res 2016;296:47–52.

48. Halberstadt AL, Chatha M, Klein AK, Wallach J, Brandt SD. Correlation between the potency of hallucinogens in the mouse head-twitch response assay and their behavioral and subjective effects in other species. Neuropharmacology 2020;167:107933.

49. Geyer MA, Light RK, Rose GJ, Petersen LR, Horwitt DD, Adams LM, et al. A characteristic effect of hallucinogens on investigatory responding in rats. Psychopharmacology (Berl.) 1979;65:35–40.

50. Adams LM, Geyer MA. A proposed animal model for hallucinogens based on LSD’s effects on patterns of exploration in rats. Behav Neurosci 1985;99;881–90.

51. Berridge KC, Aldridge JW, Houchard KR, Zhuang X. Sequential super-stereotypy of an instinctive fixed action pattern in hyper-dopaminergic mutant mice: a model of obsessive compulsive disorder and Tourette’s. BMC Biol 2005;3:4.

52. Schmid Y, Enzler F, Gasser P, Grouzmann E, Preller KH, Vollenweider FX, et al. Acute effects of lysergic acid diethylamide in healthy subjects. Biol Psychiatry 2015;78:544–53.

53. Halberstadt AL, Geyer MA. LSD but not lisuride disrupts prepulse inhibition in rats by activating the 5-HT_2A_receptor. Psychopharmacology (Berl.) 2010;208:179–89.

54. Clark WG, Clark YL. Changes in body temperature after administration of antipyretics, LSD, Δ^9^-THC, CNS depressants and stimulants, hormones, inorganic ions, gases, 2,4-DNP and miscellaneous agents. Neurosci Biobehav Rev 1981;5:1–136.

55. Forster EA, Cliffe IA, Bill DJ, Dover GM, Jones D, Reilly Y, et al. A pharmacological profile of the selective silent 5-HT_1A_receptor antagonist, WAY-100635. Eur J Pharmacol 1995;281:81–8.

56. Fox MA, French HT, LaPorte JL, Blackler AR, Murphy DL. The serotonin 5-HT_2A_receptor agonist TCB-2: a behavioral and neurophysiological analysis. Psychopharnacology (Berl) 2010;212:13–23.

57. Bohn LM, Gainetdinov RR, Sotnikova TD, Medvedev IO, Lefkowitz RJ, Dykstra LA, et al. Enhanced rewarding properties of morphine, but not cocaine in βarrestin-2 knock-out mice. J Neurosci 2003;23:10265–73.

58. Beaulieu J-M, Sotnikova T D, Marion S, Lefkowitz RJ, Gainetdinov RR, Caron MG. An Akt/β-arrestin 2/PP2A signaling complex mediates dopaminergic neurotransmission and behavior. Cell 2005;122:261–73.

59. Higgins GA, Ouagazzal AM, Grottick AJ. Influence of the 5-HT_2C_receptor antagonist SB242,084 on behaviour produced by the 5-HT_2_agonist Ro60-075 and the indirect 5-HT agonist dexfenfluramine. Br J Pharmacol 2001;133:459–66.

60. Jacobs BL, Trulson ME, and Stern WC. An animal behavior model for studying the actions of LSD and related hallucinogens. Science 1976;194:741–3

61. Braff DL, Geyer MA, Swerdlow NR. Human studies of prepulse inhibition of startle: normal subjects, patient groups, and pharmacological studies. Psychopharmacology (Berl) 2001;156:234–58.

62. Wong DF, Lever JR, Hartig PR, Dannals RF, Villemagne V, Hoffman BJ, et al. Localization of serotonin 5-HT_2_receptors in living human brain by positron emission tomography using N1-([^11^C]-methyl)-2-BR-LSD. Synapse 1987;1:393–8.

63. Salmi P, Ahlenius S. Evidence for functional interactions between 5-HT_1A_and 5-HT_2A_receptors in rat thermoregulatory mechanisms. Pharmacol Toxicol 1998;82:122–7.

64. Gurevich EV, Benovic JL, Gurevich VV. Arrestin2 and arrestin3 are differentially expressed in the rat brain during postnatal development. Neuroscience. 2002;109:421–36.

