## Supplementary Information for "LSD’s effects are differentially modulated in arrestin knockout mice"

**SUPPLEMENTARY MATERIALS**

***<u>Supplementary Materials and Methods</u>***

*Subjects*

Adult male and female WT and βArr1-KO, and WT and βArr2-KO mice were used in these experiments [1-2]. All mice had been backcrossed previously to a C57BL/6J genetic background. Heterozygotes from each strain were used to generate the respective WT and KO mice for the present experiments. The mice were housed 3-5/cage in a temperature- and humidity-controlled room on a 14:10 h (lights on at 0600 h) light-dark cycle with food and water provided *ad libitum*. All experiments were conducted with an approved protocol from the Duke University Institutional Animal Care and Use Committee.

*Drugs*

Mice were administered the vehicle or different doses of (+)-LSD-(+)-tartrate (LSD; NIDA Drug Supply Program, Bethesda, MD). The LSD was dissolved in N,N-dimethyllacetamide (DMA, final volume 0.5%; Sigma-Aldrich, St. Louis, MO) and brought to volume with 5% 2-hydroxypropoyl-β-cyclodextrin (Sigma-Aldrich) in water (Mediatech Inc., Manassas, VA). The 5-HT2AR antagonist, MDL 100907 (MDL), and the 5-HT1AR antagonist, WAY 100635 (WAY), were purchased from Tocris (Bio-Techne Corp., Minneapolis, MN) and diluted in the same vehicle. All drugs were administered (i.p.) in a 5 mL/kg volume.

*Open field activity*

Motor activities were assessed in an open field (21 x 21 x 30 cm; Omnitech Electronics, Columbus, OH) illuminated at 180 lux [3]. All behaviors in the open field were filmed. To assess 5-HT2AR-mediated responses, different doses of MDL were injected just prior to placing the mice into the open field for an examination of baseline activity over the first 30 min.

Subsequently, the animals received the vehicle or LSD and were immediately returned to the open field for 90 min. Motor activity was monitored using Fusion Integra software (Omnitech). Locomotor activity was analyzed as the distance traveled, vertical activity (rearing) was assessed by vertical beam-breaks, and stereotypical activities were examined as repetitive beam-breaks less than 1 sec in 5-min blocks or separately as cumulative baseline and post-injection LSD-stimulated activities.

*Head twitch, grooming, and retrograde walking*

These behaviors were filmed during the assessment of motor activity in the open field. The responses were scored over the first 30 min following injection of the vehicle or LSD after collection of baseline activity. All scoring was performed by blinded observers. The data are expressed as the numbers of head twitches, the duration of grooming, and the incidences of retrograde walking.

*Nose-poking responses*

Nose-pokes were monitored in a mouse 5-choice serial reaction-time apparatus (Med Associates Inc., St. Albans, VT) [4]. Each operant chamber (24 x 18.5 cm) had five LED-illuminated 1.24 $cm^2$ nose-poke apertures with infrared diodes to register nose pokes into the aperture. No food or liquid reward was available during this test. Mice were injected with the vehicle or different doses of MDL and returned to their home-cages. Thirty min later, the animals were injected with the vehicle or different doses of LSD and were placed immediately into the operant chambers. The mice remained in the chamber for 30 min and then were returned to their home cages. The data are depicted as the numbers of head pokes.

*Prepulse inhibition (PPI)*

PPI of the acoustic startle response was conducted using SR-LAB chambers (San Diego Instruments, San Diego, CA) as reported [3]. Mice were injected with vehicle or different doses of MDL or with 0.1 mg/kg haloperidol and returned to their home cages. Fifteen min later the animals were treated with the vehicle or different doses of LSD and were placed into the PPI chambers. After 10 min of habituation to a white noise background (64 dB), testing began. Each test consisted of 42 trials with 6 null trials, 18 pulse-alone trials, and 18 prepulse-pulse trials. Null trials were composed of the white noise background, pulse trials consisted of 40 ms bursts of 120 dB white-noise, and prepulse-pulse trials were comprised of 20 ms pre-pulse stimuli that were 4, 8, or 12 dB above the white-noise background (6 trials/dB), followed 100 msec later by the 120 dB pulse stimulus. Testing commenced with 10 pulse-alone trials followed by combinations of the prepulse-pulse and null trials, and it terminated with 10 pulse-alone trials. PPI responses were calculated as %PPI = [1–(pre-pulse trials/startle-only trials)]*100.

*Regulation of body temperature*

Baseline body temperatures were taken (Physitemp Instruments LLC, Clifton, NY) in the absence of the vehicle or antagonist. Immediately afterwards mice were injected with the vehicle or with different doses of MDL or WAY. Fifteen min later (time 0) their body temperatures were taken and were administered the vehicle or LSD. Subsequently, body temperatures were taken at 15, 30, 60, 120, 180, and 240 min. The data are presented as body temperatures across time.

*Radioligand binding and immunohistochemistry of 5-HT2AR*

Radioligand binding experiments on mouse brains were conducted as previously detailed using [$^3$H]-ketanserin as the radioligand [5]. The 5-HT2AR immunofluorescence was performed as described [35] with a previously validated 5-HT2AR-specific antibody [36]. Mice were intracardially perfused with phosphate-buffered saline (PBS) followed by 4% paraformaldehyde (PFA) in PBS. Brains were harvested, post-fixed overnight in 4% PFA, and dehydrated in 30% sucrose. Brains were sectioned by a cryostat at 40 µm. The brain sections were washed 3 times with 0.4% Triton X-100 in PBS (TX-100/PBS) before incubating for 1 hr with a blocking buffer (5% normal donkey serum in 0.4%TX-100/PBS). Next, they were incubated for 48 hr at 4°C with the anti-5-HT2A antibody (1:250, #RA24288; Neuromics, Edian, MN). Subsequently the sections were washed 3 times with 0.1%TX-100/PBS and then incubated for 2 hr with the secondary antibody (1:1000, donkey anti-rabbit- Alexa Fluor® 594; Jackson Immunoresearch, West Grove, PA). The sections were imaged under a 20X objective using an Olympus VS120 virtual slide microscope (Olympus, Tokyo, Japan).

*Statistics*

All statistical analyses were performed with IBM SPSS Statistics 27 programs (IBM, Chicago, IL). The data are presented as means and standard errors of the mean. No sex differences were detected across experiments so this variable was collapsed. One- or two-way ANOVA or repeated measures ANOVA (RMANOVA) were used to analyze the data and these statistics were followed with Tukey or Bonferroni *post-hoc* analyses when the main effects or interactions were significant. A $p<0.05$ was considered significant.

***References for Supplementary Materials and Methods***


1. Bohn LM, Lefkowitz RJ, Gainetdinov RR, Peppel K, Caron MG, Lin FT. Enhanced morphine analgesia in mice lacking β-arrestin 2. Science 1999;286:2495-8.
2. Kim J, Grotegut CA, Wisler JW, Li T, Mao L, Chen M, et al. β-Arrestin 1 regulates β2-adrenergic receptor-mediated skeletal muscle hypertrophy and contractility. Skelet Muscle 2018;8:39.
3. Park SM, Chen M, Schmerberg C, Dulman R, Rodriguiz RM, Caron MG, et al. Effects of β-arrestin-biased dopamine D2 receptor ligands on schizophrenia-like behavior in hypoglutamatergic mice. Neuropsychopharmacology 2016;41:704-15.
4. Velagapudi R, Subramaniyan S, Xiong C, Porkka F, Rodriguiz RM, Wetsel WC, et al. Orthopedic surgery triggers attention deficits in a delirium-like mouse model. Front Immunol 2019;10: 2675.
5. Yadav PN, Kroeze WK, Farrell MS, Roth BL. Agonist functional selectivity: 5-$HT_{2A}$ serotonin receptor antagonist differentially regulate 5-$HT_{2A}$ protein level in vivo. J Pharmacol Exp Ther 2011;339:99-105.

***Supplementary Tables***

**Supplementary Table S1:** Statistical analyses for Supplementary Figure S1a-f, β-arrestin 1: open field motor activity in 5-min segments.

*RMANOVA – Baseline Locomotion (0-30 min)*

| | Variable | Degrees of Freedom | F-Statistic | *p*-Value |
|---|---|---|---|---|
| Within Subjects | Time | 5,465 | 261.810 | <0.001 |
| | Time x Genotype | 5,465 | 0.539 | 0.747 |
| | Time x Treatment | 20,465 | 1.165 | 0.261 |
| | Time x Genotype x Treatment | 20,465 | 1.101 | 0.345 |
| Between Subjects | Genotype | 1,93 | 0.281 | 0.597 |
| | Treatment | 4,93 | 2.309 | 0.064 |
| | Genotype x Treatment | 4,93 | 0.230 | 0.921 |

*RMANOVA – Post-Injection Locomotion (31-120 min)*

| | Variable | Degrees of Freedom | F-Statistic | *p*-Value |
|---|---|---|---|---|
| Within Subjects | Time | 17,1581 | 13.894 | <0.001 |
| | Time x Genotype | 17,1581 | 0.768 | 0.732 |
| | Time x Treatment | 68,1581 | 5.806 | <0.001 |
| | Time x Genotype x Treatment | 68,1581 | 0.908 | 0.688 |
| Between Subjects | Genotype | 1,93 | 0.074 | 0.786 |
| | Treatment | 4,93 | 16.916 | <0.001 |
| | Genotype x Treatment | 4,93 | 1.188 | 0.321 |

*RMANOVA – Baseline Vertical Activity (0-30 min)*

| | Variable | Degrees of Freedom | F-Statistic | *p*-Value |
|---|---|---|---|---|
| Within Subjects | Time | 5,465 | 31.981 | <0.001 |
| | Time x Genotype | 5,465 | 1.804 | 0.111 |
| | Time x Treatment | 20,465 | 3.575 | <0.001 |
| | Time x Genotype x Treatment | 20,465 | 0.758 | 0.764 |
| Between Subjects | Genotype | 1,93 | 1.237 | 0.269 |
| | Treatment | 4,93 | 6.943 | <0.001 |
| | Genotype x Treatment | 4,93 | 1.974 | 0.105 |

**Supplementary Table S1:** Continued for β-arrestin 1 mice.

*RMANOVA – Post-Injection Vertical Activity (31-120 min)*

| | Variable | Degrees of Freedom | F-Statistic | *p*-Value |
|---|---|---|---|---|
| Within Subjects | Time | 17,1581 | 6.261 | <0.001 |
| | Time x Genotype | 17,1581 | 1.711 | 0.035 |
| | Time x Treatment | 68,1581 | 1.572 | 0.002 |
| | Time x Genotype x Treatment | 68,1581 | 0.940 | 0.617 |
| Between Subjects | Genotype | 1,93 | 1.561 | 0.215 |
| | Treatment | 4,93 | 5.108 | 0.001 |
| | Genotype x Treatment | 4,93 | 0.300 | 0.878 |

*RMANOVA – Baseline Stereotypical Activity (0-30 min)*

| | Variable | Degrees of Freedom | F-Statistic | *p*-Value |
|---|---|---|---|---|
| Within Subjects | Time | 5,465 | 172.097 | <0.001 |
| | Time x Genotype | 5,465 | 0.204 | 0.961 |
| | Time x Treatment | 20,465 | 1.751 | 0.023 |
| | Time x Genotype x Treatment | 20,465 | 1.339 | 0.149 |
| Between Subjects | Genotype | 1,93 | 1.984 | 0.162 |
| | Treatment | 4,93 | 7.110 | <0.001 |
| | Genotype x Treatment | 4,93 | 0.693 | 0.598 |

*RMANOVA – Post-Injection Stereotypical Activity (31-120 min)*

| | Variable | Degrees of Freedom | F-Statistic | *p*-Value |
|---|---|---|---|---|
| Within Subjects | Time | 17,1581 | 8.859 | <0.001 |
| | Time x Genotype | 17,1581 | 0.465 | 0.968 |
| | Time x Treatment | 68,1581 | 4.399 | <0.001 |
| | Time x Genotype x Treatment | 68,1581 | 0.828 | 0.839 |
| Between Subjects | Genotype | 1,93 | 1.679 | 0.198 |
| | Treatment | 4,93 | 5.720 | <0.001 |
| | Genotype x Treatment | 4,93 | 0.087 | 0.986 |

**Supplementary Table S2:** Statistical analyses for Supplementary Figure S2a-f, β-arrestin 2: open field motor activity in 5-min segments.

*RMANOVA – Baseline Locomotion (0-30 min)*

| | Variable | Degrees of Freedom | F-Statistic | *p*-Value |
|---|---|---|---|---|
| Within Subjects | Time | 5,480 | 143.046 | <0.001 |
| | Time x Genotype | 5,480 | 1.375 | 0.232 |
| | Time x Treatment | 25,480 | 2.117 | 0.001 |
| | Time x Genotype x Treatment | 25,480 | 0.894 | 0.614 |
| Between Subjects | Genotype | 1,96 | 1.258 | 0.265 |
| | Treatment | 5,96 | 1.205 | 0.313 |
| | Genotype x Treatment | 5,96 | 1.041 | 0.398 |

*RMANOVA – Post-Injection Locomotion (31-120 min)*

| | Variable | Degrees of Freedom | F-Statistic | *p*-Value |
|---|---|---|---|---|
| Within Subjects | Time | 17,1632 | 1.483 | 0.092 |
| | Time x Genotype | 17,1632 | 0.607 | 0.889 |
| | Time x Treatment | 85,1632 | 1.632 | <0.001 |
| | Time x Genotype x Treatment | 85,1632 | 0.821 | 0.878 |
| Between Subjects | Genotype | 1,96 | 3.641 | 0.059 |
| | Treatment | 5,96 | 18.578 | <0.001 |
| | Genotype x Treatment | 5,96 | 5.273 | <0.001 |

*RMANOVA – Baseline Vertical Activity (0-30 min)*

| | Variable | Degrees of Freedom | F-Statistic | *p*-Value |
|---|---|---|---|---|
| Within Subjects | Time | 5,480 | 27.620 | <0.001 |
| | Time x Genotype | 5,480 | 1.495 | 0.190 |
| | Time x Treatment | 25,480 | 1.506 | 0.057 |
| | Time x Genotype x Treatment | 25,480 | 0.848 | 0.680 |
| Between Subjects | Genotype | 1,96 | 0.020 | 0.889 |
| | Treatment | 5,96 | 1.769 | 0.126 |
| | Genotype x Treatment | 5,96 | 1.330 | 0.258 |

*RMANOVA – Post-Injection Vertical Activity (31-120 min)*

| | Variable | Degrees of Freedom | F-Statistic | *p*-Value |
|---|---|---|---|---|
| Within Subjects | Time | 17,1632 | 2,006 | 0.009 |
| | Time x Genotype | 17,1632 | 1.104 | 0.343 |
| | Time x Treatment | 85,1632 | 1.019 | 0.434 |
| | Time x Genotype x Treatment | 85,1632 | 1.314 | 0.032 |
| Between Subjects | Genotype | 1,96 | 0.819 | 0.368 |
| | Treatment | 5,96 | 7.150 | <0.001 |
| | Genotype x Treatment | 5,96 | 3.437 | 0.007 |

**Supplementary Table S2:** Continued for β-arrestin 2 mice.

*RMANOVA – Baseline Stereotypical Activity (0-30 min)*

| | Variable | Degrees of Freedom | F-Statistic | *p*-Value |
|---|---|---|---|---|
| Within Subjects | Time | 5,480 | 48.765 | <0.001 |
| | Time x Genotype | 5,480 | 0.604 | 0.697 |
| | Time x Treatment | 25,480 | 1.118 | 0.317 |
| | Time x Genotype x Treatment | 25,480 | 0.947 | 0.539 |
| Between Subjects | Genotype | 1,96 | 0.085 | 0.772 |
| | Treatment | 5,96 | 1.321 | 0.262 |
| | Genotype x Treatment | 5,96 | 0.480 | 0.790 |

*RMANOVA – Post-Injection Stereotypical Activity (31-120 min)*

| | Variable | Degrees of Freedom | F-Statistic | *p*-Value |
|---|---|---|---|---|
| Within Subjects | Time | 17,1632 | 1.120 | 0.327 |
| | Time x Genotype | 17,1632 | 0.669 | 0.836 |
| | Time x Treatment | 85,1632 | 1.395 | 0.012 |
| | Time x Genotype x Treatment | 85,1632 | 1.017 | 0.440 |
| Between Subjects | Genotype | 1,96 | 0.002 | 0.967 |
| | Treatment | 5,96 | 4.242 | 0.002 |
| | Genotype x Treatment | 5,96 | 1.855 | 0.109 |

**Supplementary Table S3.** Baseline distance travelled, vertical activity, and stereotypies for β-arrestin 1 mice.

| Motor Response | WT[a] | βArr1-KO[a] |
|---|---|---|
| *Distance Travelled* | | |
| Vehicle | 1033.6 ± 88.35 | 978.9 ± 93.79 |
| 0.5 mg/kg MDL - Vehicle | 853.1 ± 89.82 | 820.7 ± 29.29 |
| Vehicle + 0.3 mg/kg LSD | 984.4 ± 63.07 | 926.8 ± 59.14 |
| 0.1 mg/kg MDL + 0.3 mg/kg LSD | 831.2 ± 49.49 | 783.1 ± 96.35 |
| 0.5 mg/kg MDL + 0.3 mg/kg LSD | 819.1 ± 42.20 | 878.9 ± 107.56 |
| *Vertical Activity* | | |
| Vehicle | 393.4 ± 60.03 | 267.4 ± 69.58 |
| 0.5 mg/kg MDL - Vehicle | 211.3 ± 33.01 | 364.4 ± 72.51 |
| Vehicle + 0.3 mg/kg LSD | 363.9 ± 79.27 | 317.6 ± 41.56 |
| 0.1 mg/kg MDL + 0.3 mg/kg LSD[b] | 199.0 ± 20.55 | 113.4 ± 25.17 |
| 0.5 mg/kg MDL + 0.3 mg/kg LSD[b] | 193.0 ± 28.21 | 117.7 ± 22.31 |
| *Stereotypical Activity* | | |
| Vehicle | 4244.6 ± 379.95 | 4004.3 ± 460.29 |
| 0.5 mg/kg MDL - Vehicle | 3227.9 ± 502.66 | 3542.4 ± 221.92 |
| Vehicle + 0.3 mg/kg LSD | 4047.6 ± 328.81 | 3556.0 ± 311.30 |
| 0.1 mg/kg MDL + 0.3 mg/kg LSD[c,d] | 2643.0 ± 246.27 | 2335.9 ± 234.63 |
| 0.5 mg/kg MDL + 0.3 mg/kg LSD[c] | 3366.8 ± 164.17 | 2571.4 ± 281.79 |

[a]N = 8-17 mice/genotype/treatment.

[b]For overall rearing, the groups to receive the vehicle or LDS were significantly different from the 0.1 or 0.5 mg/kg MDL plus LSD groups.

[c]For overall stereotypical activity, the group to receive the vehicle was significantly different from the 0.1 or 0.5 mg/kg MDL plus LSD groups.

[d]For overall stereotypical activity, the group to receive LSD was significantly different from the 0.1 mg/kg MDL plus LSD group.

**Supplementary Table S4.** Statistical analyses for Figure 1a, c, and e, and Supplementary Table S3; β-arrestin 1: open field cumulative motor activity.

*ANOVA – Cumulative Baseline Locomotion (0-30 min)*

| | Variable | Degrees of Freedom | F-Statistic | *p*-Value |
|---|---|---|---|---|
| Between Subjects | Genotype | 1,93 | 0.281 | 0.597 |
| | Treatment | 4,93 | 2.309 | 0.064 |
| | Genotype x Treatment | 4,93 | 0.230 | 0.921 |

*ANOVA – Cumulative Post-Injection Locomotion (31-120 min)*

| | Variable | Degrees of Freedom | F-Statistic | *p*-Value |
|---|---|---|---|---|
| Between Subjects | Genotype | 1,93 | 0.074 | 0.786 |
| | Treatment | 4,93 | 18.916 | <0.001 |
| | Genotype x Treatment | 4,93 | 1.188 | 0.321 |

*ANOVA – Cumulative Baseline Vertical Activity (0-30 min)*

| | Variable | Degrees of Freedom | F-Statistic | *p*-Value |
|---|---|---|---|---|
| Between Subjects | Genotype | 1,93 | 1.237 | 0.269 |
| | Treatment | 4,93 | 6.943 | <0.001 |
| | Genotype x Treatment | 4,93 | 1.974 | 0.105 |

*ANOVA – Cumulative Post-Injection Vertical Activity (31-120 min)*

| | Variable | Degrees of Freedom | F-Statistic | *p*-Value |
|---|---|---|---|---|
| Between Subjects | Genotype | 1,93 | 1.561 | 0.215 |
| | Treatment | 4,93 | 5.108 | 0.001 |
| | Genotype x Treatment | 4,93 | 0.300 | 0.878 |

*ANOVA – Cumulative Baseline Stereotypical Activity (0-30 min)*

| | Variable | Degrees of Freedom | F-Statistic | *p*-Value |
|---|---|---|---|---|
| Between Subjects | Genotype | 1,93 | 1.984 | 0.162 |
| | Treatment | 4,93 | 7.110 | <0.001 |
| | Genotype x Treatment | 4,93 | 0.693 | 0.598 |

*ANOVA – Cumulative Post-Injection Stereotypical Activity (31-120 min)*

| | Variable | Degrees of Freedom | F-Statistic | *p*-Value |
|---|---|---|---|---|
| Between Subjects | Genotype | 1,93 | 1.679 | 0.199 |
| | Treatment | 4,93 | 5.720 | <0.001 |
| | Genotype x Treatment | 4,93 | 0.087 | 0.986 |

**Supplementary Table S5.** Baseline distance travelled, vertical activity, and stereotypies for β-arrestin 2 mice.

| **Motor Response** | **WT**[a] | **βArr2-KO**[a] |
|---|---|---|
| *Distance Travelled* | | |
| Vehicle | 489.34 ± 80.06 | 346.74 ± 49.35 |
| 0.5 mg/kg MDL - Vehicle | 256.78 ± 78.79 | 315.93 ± 57.03 |
| Vehicle + 0.3 mg/kg LSD | 519.58 ± 125.65 | 314.36 ± 44.71 |
| 0.05 mg/kg MDL + 0.3 mg/kg LSD | 407.60 ± 60.61 | 310.51 ± 45.67 |
| 0.15 mg/kg MDL + 0.3 mg/kg LSD | 470.92 ± 109.35 | 454.93 ± 58.01 |
| 0.5 mg/kg MDL + 0.3 mg/kg LSD | 296.28 ± 46.81 | 381.95 ± 74.91 |
| *Vertical Activity* | | |
| Vehicle | 50.80 ± 18.22 | 24.00 ± 6.35 |
| 0.5 mg/kg MDL - Vehicle | 11.63 ± 4.28 | 12.62 ± 5.07 |
| Vehicle + 0.3 mg/kg LSD | 39.41 ± 10.58 | 22.30 ± 5.39 |
| 0.05 mg/kg MDL + 0.3 mg/kg LSD | 36.12 ± 12.09 | 31.62 ± 9.53 |
| 0.1 mg/kg MDL + 0.3 mg/kg LSD | 36.00 ± 12.51 | 63.37 ± 24.12 |
| 0.5 mg/kg MDL + 0.3 mg/kg LSD | 23.12 ± 9.20 | 38.62 ± 14.32 |
| *Stereotypical Activity* | | |
| Vehicle | 1428.10 ± 178.43 | 1197.40 ± 88.91 |
| 0.5 mg/kg MDL - Vehicle | 865.37 ± 188.64 | 960.13 ± 135.06 |
| Vehicle + 0.3 mg/kg LSD | 1229.25 ± 156.81 | 1074.20 ± 95.65 |
| 0.05 mg/kg MDL + 0.3 mg/kg LSD | 1174.13 ± 115.34 | 1084.75 ± 196.94 |
| 0.1 mg/kg MDL + 0.3 mg/kg LSD | 1131.40 ± 184.69 | 1224.12 ± 155.35 |
| 0.5 mg/kg MDL + 0.3 mg/kg LSD | 1064.12 ± 194.88 | 1192.75 ± 133.49 |

[a]N = 8-12 mice/genotype/treatment.

**Supplementary Table S6:** Statistical analyses for Figure 1b, d, and f, and Supplementary Table S5; βArr2: open field cumulative motor activity.

*ANOVA – Cumulative Baseline Locomotion (0-30 min)*

| | Variable | Degrees of Freedom | F-Statistic | *p*-Value |
|---|---|---|---|---|
| Between Subjects | Genotype | 1,96 | 1.258 | 0.265 |
| | Treatment | 5,96 | 1.205 | 0.313 |
| | Genotype x Treatment | 5,96 | 1.041 | 0.398 |

*ANOVA – Cumulative Post-Injection Locomotion (31-120 min)*

| | Variable | Degrees of Freedom | F-Statistic | *p*-Value |
|---|---|---|---|---|
| Between Subjects | Genotype | 1,96 | 3.641 | 0.059 |
| | Treatment | 5,96 | 18.578 | <0.001 |
| | Genotype x Treatment | 5,96 | 5.273 | <0.001 |

*ANOVA – Cumulative Baseline Vertical Activity (0-30 min)*

| | Variable | Degrees of Freedom | F-Statistic | *p*-Value |
|---|---|---|---|---|
| Between Subjects | Genotype | 1,96 | 0.020 | 0.889 |
| | Treatment | 5,96 | 1.769 | 0.126 |
| | Genotype x Treatment | 5,96 | 1.330 | 0.258 |

*ANOVA – Cumulative Post-Injection Vertical Activity (31-120 min)*

| | Variable | Degrees of Freedom | F-Statistic | *p*-Value |
|---|---|---|---|---|
| Between Subjects | Genotype | 1,96 | 0.819 | 0.368 |
| | Treatment | 5,96 | 7.150 | <0.001 |
| | Genotype x Treatment | 5,96 | 3.437 | 0.007 |

*ANOVA – Cumulative Baseline Stereotypical Activity (0-30 min)*

| | Variable | Degrees of Freedom | F-Statistic | *p*-Value |
|---|---|---|---|---|
| Between Subjects | Genotype | 1,96 | 0.085 | 0.772 |
| | Treatment | 5,96 | 1.321 | 0.262 |
| | Genotype x Treatment | 5,96 | 0.480 | 0.790 |

*ANOVA – Cumulative Post-Injection Stereotypical Activity (31-120 min)*

| | Variable | Degrees of Freedom | F-Statistic | *p*-Value |
|---|---|---|---|---|
| Between Subjects | Genotype | 1,96 | 0.002 | 0.967 |
| | Treatment | 5,96 | 4.242 | 0.002 |
| | Genotype x Treatment | 5,96 | 1.855 | 0.109 |

**Supplementary Table S7:** Statistical analyses for Figure 2a, c, e, and g, βArr1: LSD-induced behavioral responses.

*ANOVA – Head Twitch*

| | Variable | Degrees of Freedom | F-Statistic | *p*-Value |
|---|---|---|---|---|
| Between Subjects | Genotype | 1,93 | 0.251 | 0.617 |
| | Treatment | 4,93 | 114.447 | <0.001 |
| | Genotype x Treatment | 4,93 | 0.574 | 0.682 |

*ANOVA – Grooming*

| | Variable | Degrees of Freedom | F-Statistic | *p*-Value |
|---|---|---|---|---|
| Between Subjects | Genotype | 1,93 | 2.586 | 0.111 |
| | Treatment | 4,93 | 61.232 | <0.001 |
| | Genotype x Treatment | 4,93 | 0.290 | 0.884 |

*ANOVA – Retrograde Walking*

| | Variable | Degrees of Freedom | F-Statistic | *p*-Value |
|---|---|---|---|---|
| Between Subjects | Genotype | 1,93 | 0.973 | 0.326 |
| | Treatment | 4,93 | 43.899 | <0.001 |
| | Genotype x Treatment | 4,93 | 0.106 | 0.980 |

*ANOVA – Nose Poking*

| | Variable | Degrees of Freedom | F-Statistic | *p*-Value |
|---|---|---|---|---|
| Between Subjects | Genotype | 1,89 | 0.468 | 0.496 |
| | Treatment | 4,89 | 60.656 | <0.001 |
| | Genotype x Treatment | 4,89 | 1.723 | 0.152 |

**Supplementary Table S8:** Statistical analyses for Figure 2b, d, f, and h, βArr2: LSD-induced behavioral responses.

*ANOVA – Head Twitch*

| | Variable | Degrees of Freedom | F-Statistic | *p*-Value |
|---|---|---|---|---|
| Between Subjects | Genotype | 1,96 | 31.271 | <0.001 |
| | Treatment | 5,96 | 41.567 | <0.001 |
| | Genotype x Treatment | 5,96 | 7.734 | <0.001 |

*ANOVA – Grooming*

| | Variable | Degrees of Freedom | F-Statistic | *p*-Value |
|---|---|---|---|---|
| Between Subjects | Genotype | 1,96 | 51.972 | <0.001 |
| | Treatment | 5,96 | 27.987 | <0.001 |
| | Genotype x Treatment | 5,96 | 7.953 | <0.001 |

*ANOVA – Retrograde Walking*

| | Variable | Degrees of Freedom | F-Statistic | *p*-Value |
|---|---|---|---|---|
| Between Subjects | Genotype | 1,96 | 1.418 | 0.237 |
| | Treatment | 5,96 | 13.028 | <0.001 |
| | Genotype x Treatment | 5,96 | 5.199 | <0.001 |

*ANOVA – Nose Poking*

| | Variable | Degrees of Freedom | F-Statistic | *p*-Value |
|---|---|---|---|---|
| Between Subjects | Genotype | 1,125 | 2.160 | 0.144 |
| | Treatment | 5,125 | 7.512 | <0.001 |
| | Genotype x Treatment | 5,125 | 4.769 | 0.001 |

**Supplementary Table S9:** Statistical analyses for Figure 3a, βArr1: Prepulse Inhibition – MDL and LSD, and for Supplementary Figure S3a-b for null and pulse activities.

*ANOVA – Null Activity*

| | Variable | Degrees of Freedom | F-Statistic | *p*-Value |
|---|---|---|---|---|
| Between Subjects | Genotype | 1,91 | 0.725 | 0.397 |
| | Treatment | 4,91 | 1.099 | 0.362 |
| | Genotype x Treatment | 4,91 | 0.319 | 0.864 |

*ANOVA – Startle Activity*

| | Variable | Degrees of Freedom | F-Statistic | *p*-Value |
|---|---|---|---|---|
| Between Subjects | Genotype | 1,91 | 1.680 | 0.198 |
| | Treatment | 4,91 | 0.409 | 0.802 |
| | Genotype x Treatment | 4,91 | 0.333 | 0.855 |

*RMANOVA -- PPI*

| | Variable | Degrees of Freedom | F-Statistic | *p*-Value |
|---|---|---|---|---|
| Within Subjects | PPI | 1,91 | 487.507 | <0.001 |
| | PPI x Genotype | 1,91 | 9.162 | 0.003 |
| | PPI x Treatment | 4,91 | 7.944 | <0.001 |
| | PPI x Genotype x Treatment | 4,91 | 2.611 | 0.041 |
| Between Subjects | Genotype | 1,91 | 25.358 | <0.001 |
| | Treatment | 4,91 | 11.435 | <0.001 |
| | Genotype x Treatment | 4,91 | 2.394 | 0.052 |

**Supplementary Table S10:** Statistical analyses for Figure 3b, βArr1: prepulse inhibition – haloperidol and LSD, and for Supplementary Figure S3c-d for null and pulse activities**.**

*ANOVA – Null Activity*

| | Variable | Degrees of Freedom | F-Statistic | *p*-Value |
|---|---|---|---|---|
| Between Subjects | Genotype | 1,72 | 0.132 | 0.717 |
| | Treatment | 3,72 | 5.673 | 0.002 |
| | Genotype x Treatment | 3,72 | 0.255 | 0.858 |

*ANOVA – Startle Activity*

| | Variable | Degrees of Freedom | F-Statistic | *p*-Value |
|---|---|---|---|---|
| Between Subjects | Genotype | 1,72 | 5.015 | 0.028 |
| | Treatment | 3,72 | 0.838 | 0.477 |
| | Genotype x Treatment | 3,72 | 2.522 | 0.064 |

*RMANOVA -- PPI*

| | Variable | Degrees of Freedom | F-Statistic | *p*-Value |
|---|---|---|---|---|
| Within Subjects | PPI | 2,144 | 233.984 | <0.001 |
| | PPI x Genotype | 2,144 | 1.953 | 0.146 |
| | PPI x Treatment | 6,144 | 4.566 | <0.001 |
| | PPI x Genotype x Treatment | 6,144 | 0.671 | 0.673 |
| Between Subjects | Genotype | 1,72 | 7.563 | 0.008 |
| | Treatment | 3,72 | 9.591 | <0.001 |
| | Genotype x Treatment | 3,72 | 0.525 | 0.667 |

**Supplementary Table S11:** Statistical analyses for Figure 3c, βArr2: prepulse inhibition – MDL and LSD, and for Supplementary Figure S3e-f for null and pulse activities.

*ANOVA – Null Activity*

| | Variable | Degrees of Freedom | F-Statistic | *p*-Value |
|---|---|---|---|---|
| Between Subjects | Genotype | 1,74 | 0.035 | 0.852 |
| | Treatment | 4,74 | 5.439 | 0.001 |
| | Genotype x Treatment | 4,74 | 0.705 | 0.591 |

*ANOVA – Startle Activity*

| | Variable | Degrees of Freedom | F-Statistic | *p*-Value |
|---|---|---|---|---|
| Between Subjects | Genotype | 1,74 | 0.126 | 0.724 |
| | Treatment | 4,74 | 2.321 | 0.065 |
| | Genotype x Treatment | 4,74 | 0.448 | 0.773 |

*RMANOVA -- PPI*

| | Variable | Degrees of Freedom | F-Statistic | *p*-Value |
|---|---|---|---|---|
| Within Subjects | PPI | 1,74 | 580.044 | <0.001 |
| | PPI x Genotype | 1,74 | 2.321 | 0.132 |
| | PPI x Treatment | 4,74 | 0.668 | 0.616 |
| | PPI x Genotype x Treatment | 4,74 | 1.331 | 0.267 |
| Between Subjects | Genotype | 1,74 | 18.823 | <0.001 |
| | Treatment | 4,74 | 3.953 | 0.006 |
| | Genotype x Treatment | 4,74 | 5.660 | <0.001 |

**Supplementary Table S12:** Statistical analyses for Figure 4a-h βArr1: temperature regulation – MDL, WAY, and LSD.

*Figure 4a-b: RMANOVA – Vehicle, MDL100907, and LSD*

| | Variable | Degrees of Freedom | F-Statistic | *p*-Value |
|---|---|---|---|---|
| Within Subjects | Time | 6,354 | 24.307 | <0.001 |
| | Time x Genotype | 6,354 | 12.565 | <0.001 |
| | Time x Treatment | 12,354 | 18.053 | <0.001 |
| | Time x Genotype x Treatment | 12,354 | 3.730 | <0.001 |
| Between Subjects | Genotype | 1,59 | 8.627 | 0.005 |
| | Treatment | 2,59 | 8.390 | 0.001 |
| | Genotype x Treatment | 2,59 | 3.216 | 0.047 |

*Figure 4c-d: RMANOVA – Vehicle, LSD, and MDL + LSD*

| | Variable | Degrees of Freedom | F-Statistic | *p*-Value |
|---|---|---|---|---|
| Within Subjects | Time | 6,462 | 40.096 | <0.001 |
| | Time x Genotype | 6,462 | 15.310 | <0.001 |
| | Time x Treatment | 18,462 | 29.740 | <0.001 |
| | Time x Genotype x Treatment | 18,462 | 4.047 | <0.001 |
| Between Subjects | Genotype | 1,77 | 7.570 | 0.007 |
| | Treatment | 3,77 | 22.323 | <0.001 |
| | Genotype x Treatment | 3,77 | 3.843 | 0.013 |

*Figure 4e-f: RMANOVA – Vehicle, LSD, and WAY100635*

| | Variable | Degrees of Freedom | F-Statistic | *p*-Value |
|---|---|---|---|---|
| Within Subjects | Time | 6,342 | 16.757 | <0.001 |
| | Time x Genotype | 6,342 | 3.480 | 0.002 |
| | Time x Treatment | 12,342 | 27.160 | <0.001 |
| | Time x Genotype x Treatment | 12,342 | 6.162 | <0.001 |
| Between Subjects | Genotype | 1,57 | 17.091 | <0.001 |
| | Treatment | 2,57 | 4.426 | 0.016 |
| | Genotype x Treatment | 2,57 | 3.795 | 0.028 |

*Figure 4g-h: RMANOVA – Vehicle, LSD, and WAY + LSD*

| | Variable | Degrees of Freedom | F-Statistic | *p*-Value |
|---|---|---|---|---|
| Within Subjects | Time | 6,450 | 28.906 | <0.001 |
| | Time x Genotype | 6,450 | 4.647 | <0.001 |
| | Time x Treatment | 18,450 | 19.125 | <0.001 |
| | Time x Genotype x Treatment | 18,450 | 4.864 | <0.001 |
| Between Subjects | Genotype | 1,75 | 13.519 | <0.001 |
| | Treatment | 3,75 | 3.669 | 0.016 |
| | Genotype x Treatment | 3,75 | 3.353 | 0.023 |

**Supplementary Table S13:** Statistical analyses for Figure 5a-h βArr2: temperature regulation – MDL, WAY, and LSD.

*Figure 5a-b: RMANOVA – Vehicle, MDL100907, and LSD*

| | Variable | Degrees of Freedom | F-Statistic | *p*-Value |
|---|---|---|---|---|
| Within Subjects | Time | 6,282 | 17.413 | <0.001 |
| | Time x Genotype | 6,282 | 3.732 | 0.001 |
| | Time x Treatment | 12,282 | 3.177 | <0.001 |
| | Time x Genotype x Treatment | 12,282 | 7.353 | <0.001 |
| Between Subjects | Genotype | 1,47 | 2.080 | 0.156 |
| | Treatment | 2,47 | 12.705 | <0.001 |
| | Genotype x Treatment | 2,47 | 0.035 | 0.965 |

*Figure 5c-d: RMANOVA – Vehicle, LSD, and MDL + LSD*

| | Variable | Degrees of Freedom | F-Statistic | *p*-Value |
|---|---|---|---|---|
| Within Subjects | Time | 6,390 | 18.009 | <0.001 |
| | Time x Genotype | 6,390 | 5.963 | <0.001 |
| | Time x Treatment | 18,390 | 3.312 | <0.001 |
| | Time x Genotype x Treatment | 18,390 | 5,230 | <0.001 |
| Between Subjects | Genotype | 1,85 | 2.451 | 0.122 |
| | Treatment | 3,85 | 11.677 | <0.001 |
| | Genotype x Treatment | 3,85 | 0.189 | 0.904 |

*Figure 5e-f: RMANOVA – Vehicle, LSD, and WAY100635D*

| | Variable | Degrees of Freedom | F-Statistic | *p*-Value |
|---|---|---|---|---|
| Within Subjects | Time | 6,342 | 8.708 | <0.001 |
| | Time x Genotype | 6,342 | 4.394 | <0.001 |
| | Time x Treatment | 12,342 | 7.231 | <0.001 |
| | Time x Genotype x Treatment | 12,342 | 7.888 | <0.001 |
| Between Subjects | Genotype | 1,57 | 7.340 | 0.009 |
| | Treatment | 2,57 | 1.560 | 0.219 |
| | Genotype x Treatment | 2,57 | 0.799 | 0.455 |

*Figure 5g-h: RMANOVA – Vehicle, LSD, and WAY + LSD*

| | Variable | Degrees of Freedom | F-Statistic | *p*-Value |
|---|---|---|---|---|
| Within Subjects | Time | 6,420 | 12.068 | <0.001 |
| | Time x Genotype | 6,420 | 10.002 | <0.001 |
| | Time x Treatment | 18,420 | 5.308 | <0.001 |
| | Time x Genotype x Treatment | 18,420 | 6.539 | <0.001 |
| Between Subjects | Genotype | 1,70 | 0.031 | 0.861 |
| | Treatment | 3,70 | 1.155 | 0.333 |
| | Genotype x Treatment | 3,70 | 1.112 | 0.350 |

**Supplementary Table S14:** Statistical analyses for Figure 6a,b: radioligand binding with samples from βArr1 and βArr2 brain**.**

*Figure 6a: ANOVA – Binding Affinity*

| | Variable | Degrees of Freedom | F-Statistic | *p*-Value |
|---|---|---|---|---|
| Between Subjects | Genotype | 3,14 | 6.855 | 0.005 |

*Figure 6b: ANOVA – $B_{max}$ Receptor Sites*

| | Variable | Degrees of Freedom | F-Statistic | *p*-Value |
|---|---|---|---|---|
| Between Subjects | Genotype | 3,14 | 6.527 | 0.006 |

***Supplementary Figures***

**Fig. S1 Effects of LSD and MDL100907 on motor activities in 5-min segments for β-arrestin 1 mice.**

A description of the experimental design can be found in the legend for Figure 1. The data are presented in 5-min intervals and the statistics are available in Supplementary Table S1. **a,b** Locomotor activities in WT and βArr1-KO mice. **c,d** Rearing activities in βArr1 animals. **e,f** Stereotypical activities in βArr1 subjects. N = 8-17 mice/group.

**Fig. S2 Effects of LSD and MDL100907 on motor activities in 5-min segments for β-arrestin 2 mice.**

The experimental procedure is described in the legend for Figure 1. Motor activities are displayed in 5-min intervals and the statistics are in Supplementary Table S2. **a,b** Locomotor activities in WT and βArr2-KO mice. **c,d** Rearing activities in βArr2 animals. **e,f** Stereotypical activities in βArr2 subjects. N = 8-12 mice/group.

**Fig. S3 Effects of LSD and MDL100907 on null and startle activities for prepulse inhibition in β-arrestin 1 and β-arrestin 2 mice.**

Mice were injected initially with MDL100907, haloperidol, or the vehicle and administered subsequently the vehicle or LSD prior to testing PPI. The statistics for the βArr1 and βArr2 responses are presentred in Supplementary Tables S9-S11. **a,b** Null and startle activities in WT and βArr1-KO mice treated with MDL and LSD. **c,d** Null and startle activities in WT and βArr1-KO mice treated with haloperidol and LSD. **e,f** Null and startle activities in WT and βArr2-KO mice treated with MDL and LSD. N = 8-12 mice/group. +$p$<0.05, vs. 0.1 mg/kg HAL + LSD overall (βArr1); ‡$p$<0.05, vs. 0.1 mg/kg MDL + LSD overall (βArr2).

**Fig. S4  Radioligand binding and immunohistochemistry of 5-HT2ARs in βArr1 and βArr2 mice.**

The statistics for the radioligand binding are located in Supplementary Table S14.  **a** Binding affinity as $K_d$ (nM) for [$^3$H]-ketanserin to membranes from βArr1 and βArr2 brains.  **b** Bmax in pmol/mg from brains of WT and βArr1-WT, and WT and βArr2-KO mice.  N = 3-5 mice/genotype.  *$p$<0.05, vs. the βArr2-KO samples.  **c-f** Representative 5-HT2AR immunofluorescence in coronal brain sections from respective βArr1 WT and KO mice, and βArr2 WT and KO mice (N = 3 mice/genotype).

***Supplementary Video Clips***

**Movie 1  Vehicle-treated WT mouse.**

Representative example of a vehicle-treated WT mouse displaying a full grooming sequence. Here. the mouse grooms it face, left flank, right flank, base of tail, and feet.  The vehicle-treated βArr1-KO and βArr2-KO show similar responses.

**Movie 2  LSD-treated WT mouse.**

Representative example of a WT mouse given LSD plus the vehicle that displays a prolonged focal grooming bout.  The mouse focuses on grooming its tail.  Other WT mice may focus on grooming the flanks or feet, rarely the face.  If the face is groomed, grooming is typically aborted soon after.

**Movie 3  LSD-treated βArr1-KO mouse.**

Representative example of a βArr1-KO mouse administered LSD plus the vehicle that shows disorganized grooming.  These mutants typically begin a grooming sequence, focusing on the face and abdomen, where they switch back and forth, before proceeding to the flanks or tail and then they rapidly repeat this sequence.

**Movie 4  LSD-treated βArr2-KO mouse.**

Representative example of a βArr2-KO mouse injected with LSD plus the vehicle.  The organization of the grooming sequence in these mice is complete and is rarely abbreviated or disrupted.

**Movie 5  MDL-treated WT mouse.**

Representative example of a WT mouse that received MDL plus the vehicle.  The organization of the grooming sequence in the WT and βArr1-KO mice is intact.

**Movie 6  MDL-treated βArr2-KO mouse.**

Representative example of a βArr2-KO mouse that received MDL plus the vehicle. While these mutants display an intact grooming sequence when treated with MDL, they also pause in the grooming bout and exhibit twitching spasticity of the muscles along the back and neck area.

**Movie 7  WT mouse treated with MDL plus LSD.**

Representative example of a WT mouse administered MDL followed by LSD. The MDL appears to restore the organization of grooming in these mice.

**Movie 8  βArr1-KO mouse treated with MDL plus LSD.**

Representative example of a βArr1-KO mouse given MDL followed by LSD. These mutants begin the sequence of grooming, but have some focal grooming on the face and abdomen, changing between these two areas before moving to the flank and completing the sequence. Note, this pattern of behavior is similar to the LSD plus vehicle sequence, except the mice do not repeat the sequence immediately upon completion.

**Movie 9  βArr2-KO mouse treated with MDL plus LSD.**

Representative example of a βArr2-KO mouse injected with MDL followed by LSD. The organization of grooming is relatively normal except there is some focal grooming mid-point and the sequence is not finished beyond the side groom. Hence, MDL given with LSD partially disturbs completion of the grooming sequence in the βArr2-KO mice.

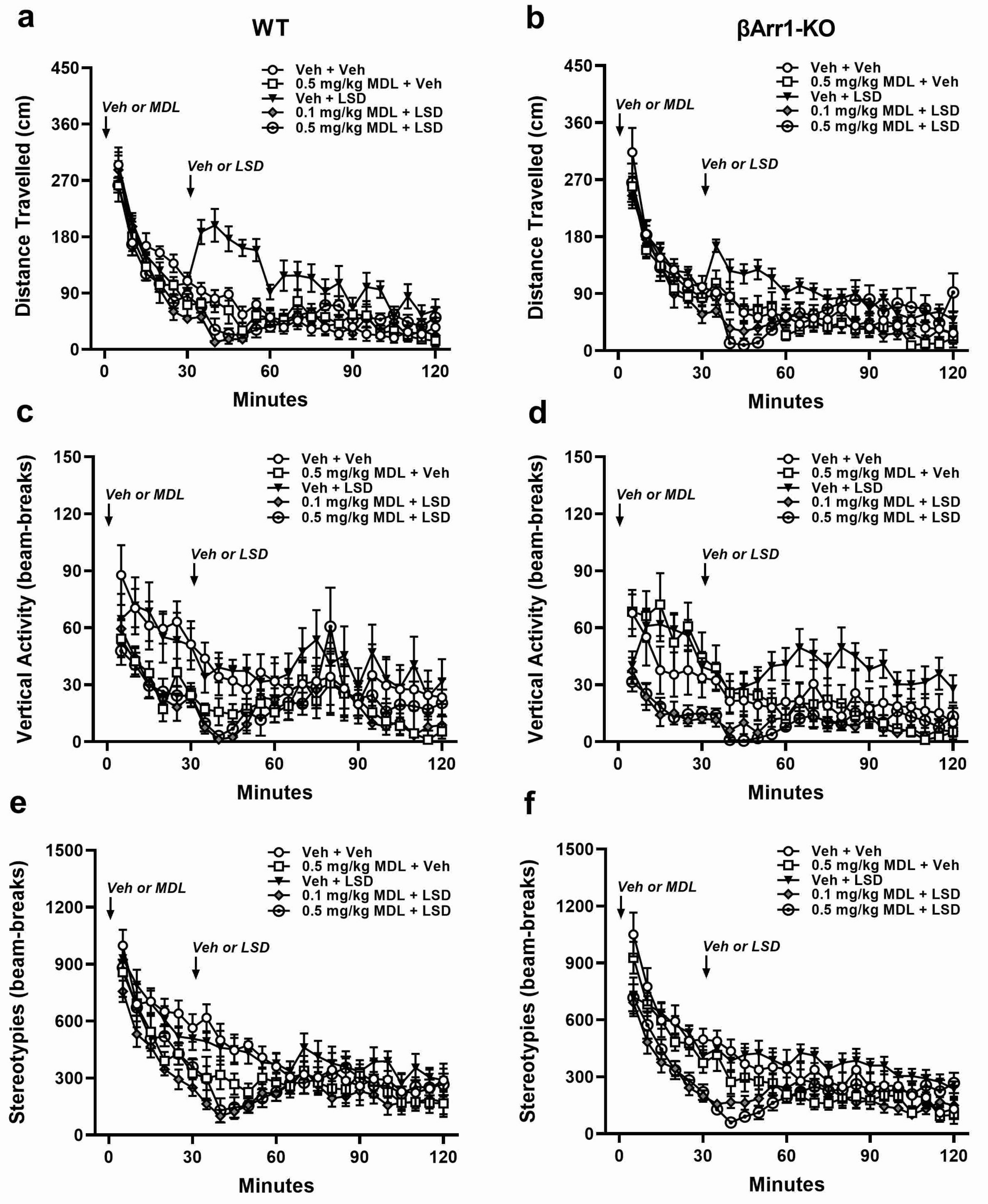

a
WT
b
βArr1-KO
Distance Travelled (cm)
Vertical Activity (beam-breaks)
Stereotypies (beam-breaks)
Minutes
Veh or MDL
Veh or LSD
Veh + Veh
0.5 mg/kg MDL + Veh
Veh + LSD
0.1 mg/kg MDL + LSD
0.5 mg/kg MDL + LSD
c
d
e
f

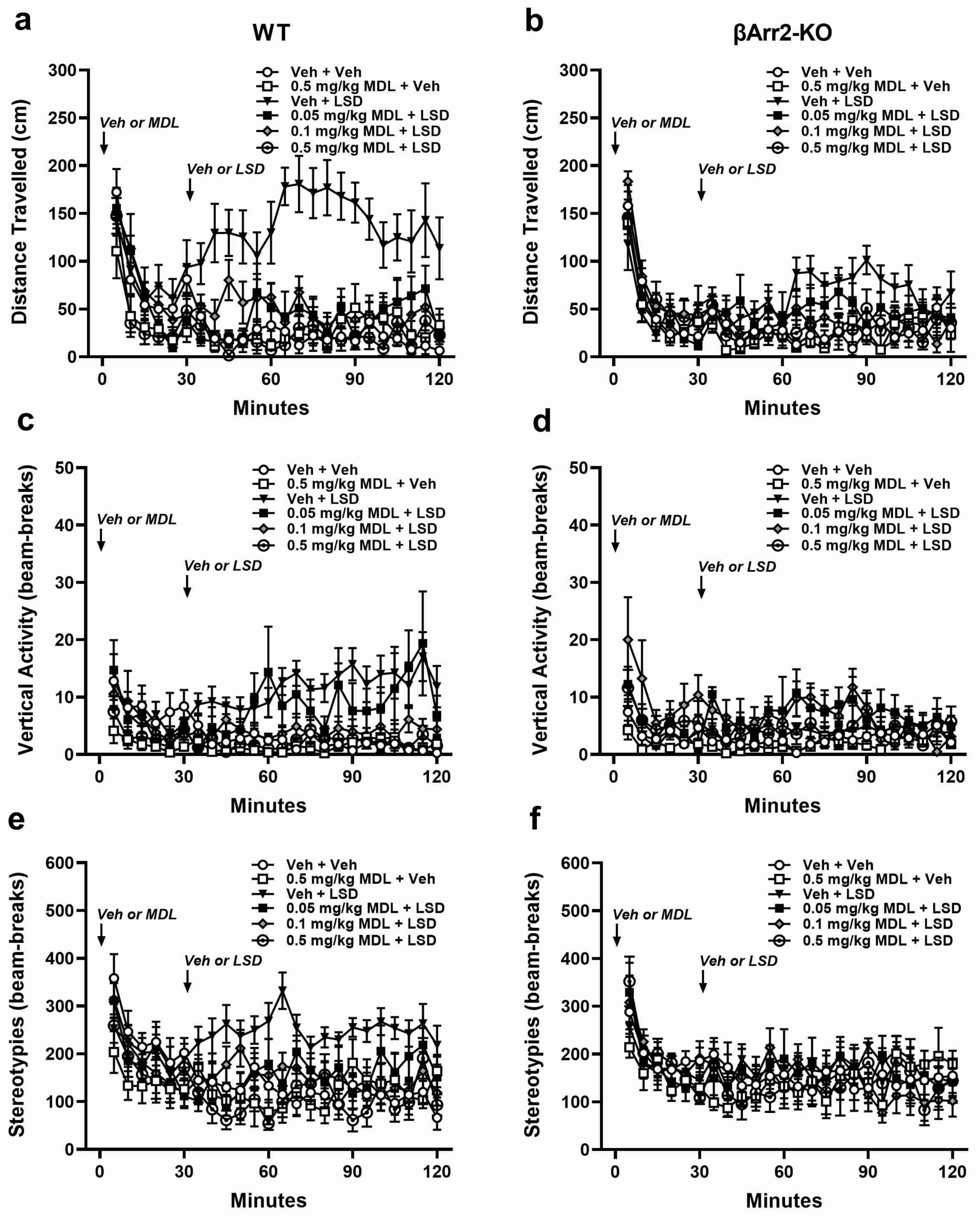
a
WT
b
βArr2-KO
Distance Travelled (cm)
Vertical Activity (beam-breaks)
Stereotypies (beam-breaks)
Minutes
Veh or MDL
Veh or LSD
Veh + Veh
0.5 mg/kg MDL + Veh
Veh + LSD
0.05 mg/kg MDL + LSD
0.1 mg/kg MDL + LSD
0.5 mg/kg MDL + LSD
c
d
e
f

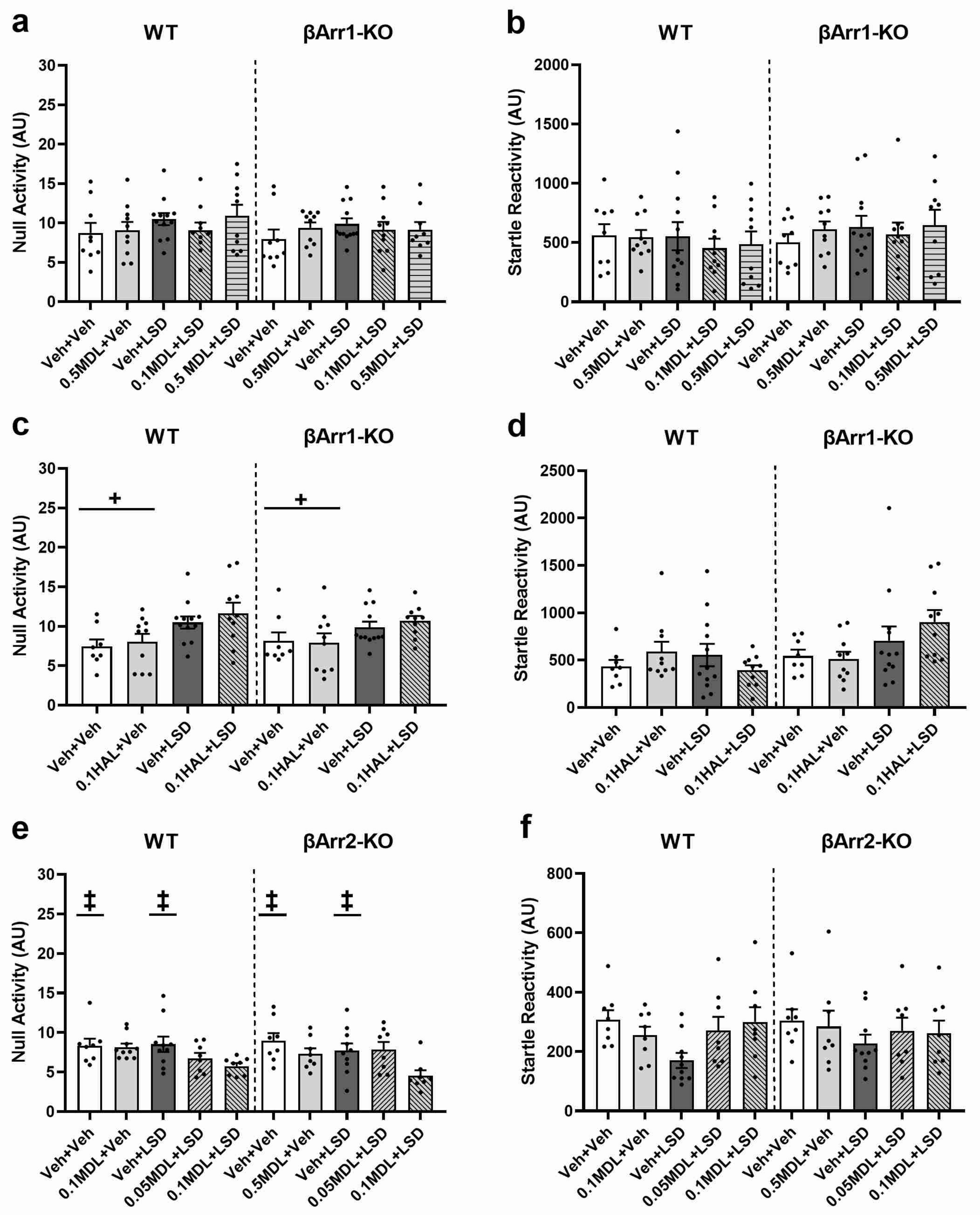
a
WT
βArr1-KO
Null Activity (AU)
Veh+Veh
0.5MDL+Veh
Veh+LSD
0.1MDL+LSD
0.5 MDL+LSD
Veh+Veh
0.5MDL+Veh
Veh+LSD
0.1MDL+LSD
0.5MDL+LSD
b
WT
βArr1-KO
Startle Reactivity (AU)
Veh+Veh
0.5MDL+Veh
Veh+LSD
0.1MDL+LSD
0.5MDL+LSD
Veh+Veh
0.5MDL+Veh
Veh+LSD
0.1MDL+LSD
0.5MDL+LSD
c
WT
βArr1-KO
+
+
Null Activity (AU)
Veh+Veh
0.1HAL+Veh
Veh+LSD
0.1HAL+LSD
Veh+Veh
0.1HAL+Veh
Veh+LSD
0.1HAL+LSD
d
WT
βArr1-KO
Startle Reactivity (AU)
Veh+Veh
0.1HAL+Veh
Veh+LSD
0.1HAL+LSD
Veh+Veh
0.1HAL+Veh
Veh+LSD
0.1HAL+LSD
e
WT
βArr2-KO
‡
‡
‡
‡
Null Activity (AU)
Veh+Veh
0.1MDL+Veh
Veh+LSD
0.05MDL+LSD
0.1MDL+LSD
Veh+Veh
0.5MDL+Veh
Veh+LSD
0.05MDL+LSD
0.1MDL+LSD
f
WT
βArr2-KO
Startle Reactivity (AU)
Veh+Veh
0.1MDL+Veh
Veh+LSD
0.05MDL+LSD
0.1MDL+LSD
Veh+Veh
0.5MDL+Veh
Veh+LSD
0.05MDL+LSD
0.1MDL+LSD

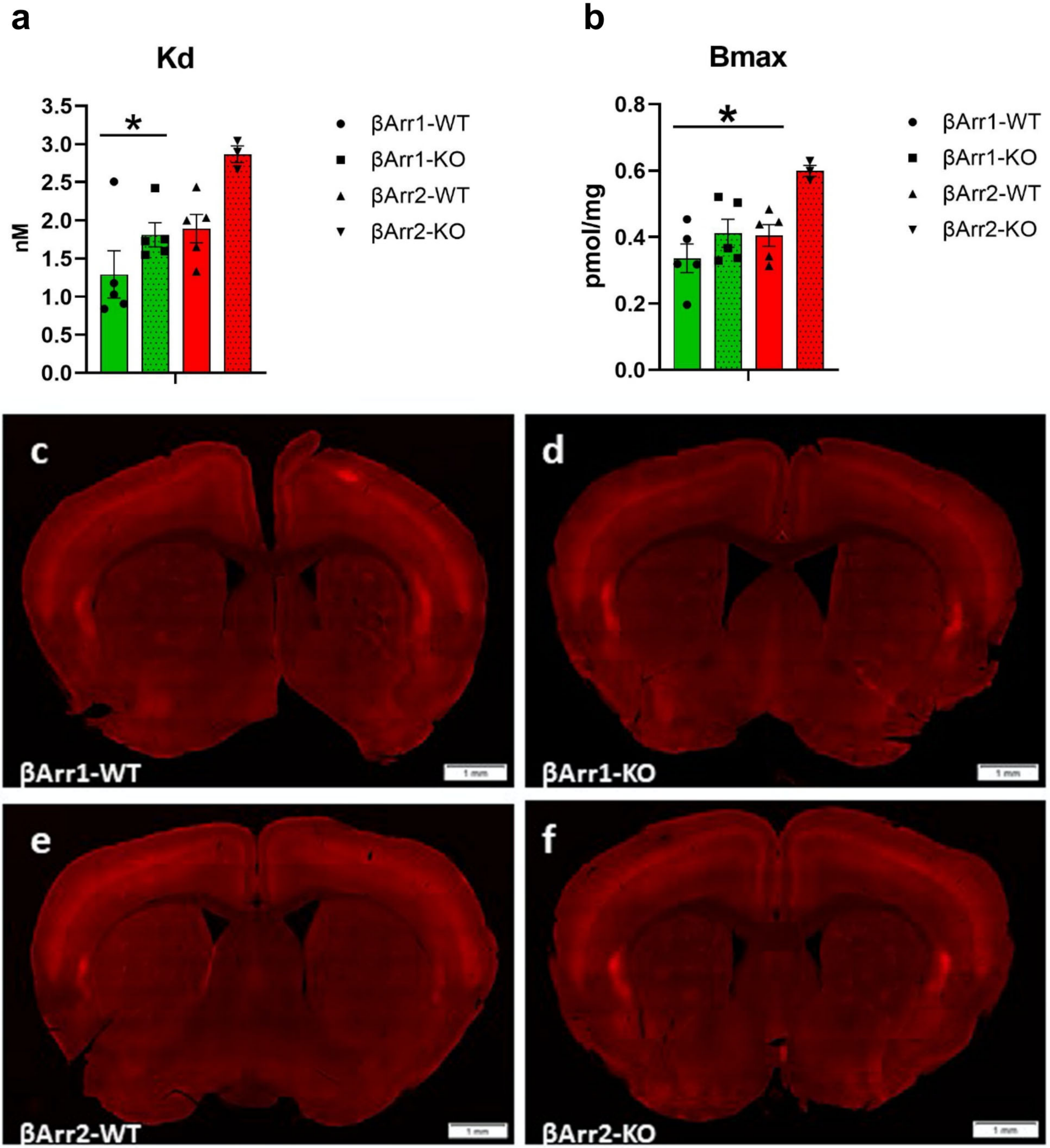

a
Kd
nM
3.5
3.0
2.5
2.0
1.5
1.0
0.5
0.0
*
βArr1-WT
βArr1-KO
βArr2-WT
βArr2-KO
b
Bmax
pmol/mg
0.8
0.6
0.4
0.2
0.0
*
βArr1-WT
βArr1-KO
βArr2-WT
βArr2-KO
c
βArr1-WT
d
βArr1-KO
e
βArr2-WT
f
βArr2-KO
